# Wildlife provisioning selects for higher pathogen virulence in hosts with incomplete immunity

**DOI:** 10.1101/2024.09.05.611527

**Authors:** Jason Cosens Walsman, Arietta E Fleming-Davies, Richard Hall, Dana Hawley

**Affiliations:** University of Pittsburgh; University of San Diego; University of Georgia; Virginia Tech

## Abstract

Anthropogenic food provisioning provides massive inputs of food to wildlife, with profound ecological and evolutionary consequences. By altering wildlife condition, density, and behavior, provisioning can influence transmission of infectious diseases and thus may impose strong selection pressure on wildlife pathogens. But surprisingly, we lack theory on the eco-evolutionary consequences of provisioning for host-pathogen dynamics. Here we develop a mathematical model of the eco-evolutionary dynamics of a wildlife pathogen under provisioning, motivated by *Mycoplasma gallisepticum*, a bacterial pathogen that emerged, spread, and changed its virulence in provisioned house finches. We model how provisioning influences the evolution of pathogen virulence, defined here as mortality associated with infection. Consistent with past empirical work, house finches recover from infection and acquire incomplete immunity; this incomplete immunity is stronger if their initial infection was with a more virulent pathogen strain. We find that, even though provisioning improves body condition, it should still select for higher virulence, and thus may actually lead to declines in host populations. These negative effects arise because provisioning magnifies the impact of incomplete immunity, selecting for higher virulence and driving host populations down. Our results highlight that food provisioning can select for more virulent pathogens, with potentially far-reaching implications for conservation.

## Introduction

Anthropogenic food subsidies to wildlife, ranging from landfills to feeding for conservation, differ from natural food sources in their quantity, quality, and spatiotemporal predictability, with profound impacts on wildlife (Oro et al. 2013). Such provisioning may have direct positive or negative impacts on the health of individual animals (Murray et al. 2016; Strandin et al. 2018); high-quality food provisioning can increase reproduction, survival and body condition, while poor-quality food or incidental exposure to contaminants in anthropogenic foods could reduce these components of fitness (Murray et al. 2016; Strandin et al. 2018). By aggregating many animals around point food sources, provisioning may also increase the transmission and spread of infectious diseases of wildlife, an area of growing concern (Becker and Hall 2014; Becker et al. 2018a; Becker et al. 2018b; Erazo et al. 2022). Regular provisioning can also have evolutionary implications, such as the evolution of increased bill length in association with backyard bird-feeding in European Great Tits (*Parus major*) (Bosse et al. 2017). Although food subsidy effects, such as loss of long-distance movements or aggregation at sites with high transmission risk (Murray et al. 2016), alter the selective environment for wildlife pathogens, to date, surprisingly little research has explored the implications of food subsidy for pathogen evolution (but see Hite and Cressler 2018; Hite and Cressler 2019). Pathogen evolution in response to food subsidy is of crucial concern, because selection on transmissibility or virulence could have negative population impacts on provisioned wildlife, and potentially favour the emergence of pathogens in novel hosts, including domestic animals and people (Plowright et al. 2017).

Infectious agents often have the capacity to evolve rapidly in response to changes in their host or environment, in ways that influence the harm they cause their hosts (i.e., their virulence) (Bonneaud et al. 2018; Cressler et al. 2016). Less virulent pathogen strains benefit by keeping their infected hosts alive and infectious for longer, while more virulent strains often exploit their hosts more rapidly, benefitting by replicating and/or transmitting more quickly (Acevedo et al. 2019). Ecological forces, such as fluctuating food availability, shift the costs and benefits of higher virulence, leading to pathogen evolution (Lopez-Pascua et al. 2014; Walsman et al. 2022). Periods of resource scarcity that influence individual host resource intake or population density can select for higher or lower virulence depending on how the immune system and pathogens compete for resources (Bosse et al. 2017; Hite and Cressler 2019). By increasing the pool of resources available for within-host pathogen replication and host immune defense, and increasing host population densities that facilitate between-host transmission, food provisioning may alter these tradeoffs with potentially divergent outcomes for pathogen fitness and evolution (Murray et al. 2016).

Whether food provisioning selects for increased pathogen virulence may depend critically on the extent to which hosts acquire immune memory responses characterized by pathogen resistance and/or tolerance from initial pathogen infection. Following initial infection, some hosts acquire incomplete immunity, that increases their resistance to reinfection, i.e., some host defense that improves host fitness but lowers pathogen fitness (e.g., lowering the rate of reinfection). If reinfections require high pathogen transmissibility, this could select for increased pathogen transmissibility and virulence. Alternatively or in addition, initial infection could prime host tolerance of reinfection, lowering the lethality of reinfection without reducing pathogen fitness (Nahrendorf et al. 2021) and altering constraints on virulence evolution. Mathematical models have provided powerful tools for parsing out the importance of host density-mediated versus defense-mediated effects of food subsidy in driving pathogen dynamics and impacts, but these models have typically ignored pathogen evolution and only explored immune-mediated increases in host resistance to, or clearance of, infection (Becker and Hall 2014; Erazo et al. 2022). Understanding the consequences of food subsidy for pathogen virulence evolution, and outcomes for host populations, requires combining evolutionary models with more complex infection epidemiologies.

Here we present a modeling framework for studying the evolution of virulence in food-subsidized wildlife hosts that integrates individual and population-level host processes with pathogen evolutionary dynamics. We center our exploration of provisioning, acquired immunity, and virulence evolution within a highly-relevant biological system. Backyard bird-feeding is a globally popular form of food subsidy that has been linked to the emergence and spread of infectious diseases for multiple songbird species (Galbraith et al. 2017; Lawson et al. 2012; Robinson et al. 2010; Tizard 2004; Wilcoxen et al. 2015), including *Mycoplasma gallisepticum* in house finches (*Haemorhous mexicanus*) (Adelman et al. 2015). Following its emergence in the eastern United States, the pathogen spread across the continent, and rapidly evolved higher virulence (Bonneaud et al. 2018; Hawley et al. 2013). Virulence is linked to a higher pathogen replication rate, increasing transmissibility. House finches can acquire incomplete immunity upon recovery from infection, and this immune priming is stronger when the pathogen strain of the initial infection was more virulent (Fleming-Davies et al. 2018). We extend an empirically parameterized model of the house-finch-Myscoplasma interaction (Fleming-Davies et al. 2018) to derive analytical conditions for the invasion of a mutant pathogen strain, and solve our model numerically to characterize how various ecological and immunological effects of provisioning should alter selection on pathogen virulence, and thus eco-evolutionary outcomes for host populations.

## Methods

### Model structure

We model the effects of provisioning on virulence evolution in the house finch-*Mycoplasma* system with variable, incomplete, acquired immunity. We follow the model of Fleming-Davies et al. 2018 (Fleming-Davies et al. 2018) using their empirically-derived parameter values as our baseline values (Eq. 1 for a single, resident strain, *r*, color-coded to match across equations and the diagram in Box 1; see Table A1 for parameter values):

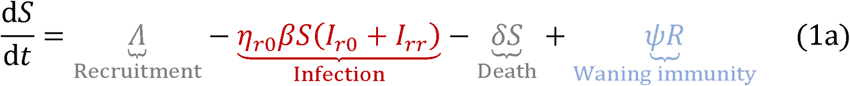

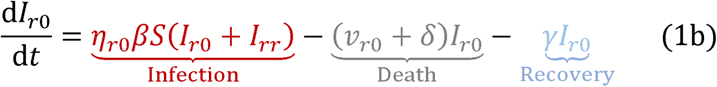

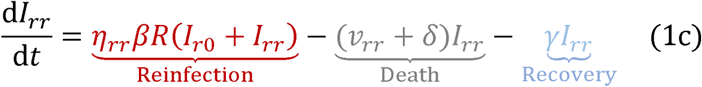

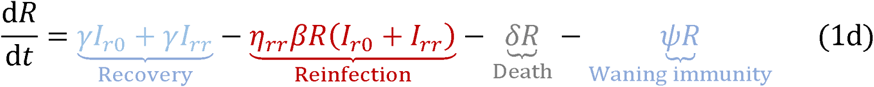

Susceptible hosts (density *S,* Eq. 1a) are produced at a fixed rate (Λ) and become infected through contact with pathogens shed by newly-infected (i.e., hosts experiencing their first infection, *I*_r0_) or reinfected hosts (*I*_rr_) at rate β (henceforth called the transmission rate) with probability η_r0_(henceforth susceptibility). Additionally, susceptible hosts can die at per-capita rate δ or arise from recovered hosts whose immunity has waned at per-capita rate ψ. Newly-infected hosts (Eq. 1b) die at an increased rate due to pathogen virulence (*v*_r0_) and recover at a per-capita rate γ, into a partially immune recovered class (*R*). Reinfected hosts (Eq. 1c) follow very similar dynamics except that they arise from infection of recovered hosts with susceptibility η*_rr_* and experience a different virulence, *v_rr_* but we assume that both infected and reinfected hosts have the same infectiousness (and thus the same transmission rate β). Recovered hosts (Eq. 1d) have acquired incomplete immunity and may become reinfected, die, or lose acquired immunity due to waning over time. Susceptibility to infection (η) and virulence (*v*) depend both on the strain of the previous infection and the strain of the current infection.

We consider pathogen evolution by modelling the invasion of a rare mutant pathogen strain, *m*. Because the frequency of *m* is very low, we may neglect the density of hosts recovered from strain *m* and therefore also the density of hosts that recovered from *m* and then became reinfected (i.e., any mutant fitness terms involving *I_mm_* can be neglected as zero). We consider the growth rates of hosts infected with strain *m* or reinfected with strain *m* after initial infection with the resident strain, *r*, at the equilibrium set by the resident strain (e.g., *S*\* *_r_*, *R*\* which affect Eqs. 2a, b):

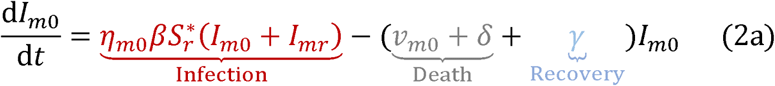

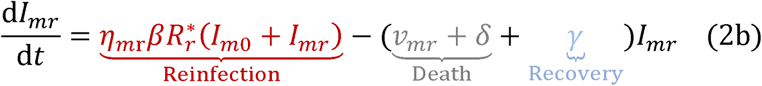

Eqs. 2a, b follow Eqs. 1b, c very closely except we group the death and recovery terms as these have equivalent implications for pathogen fitness (though not host fitness), which is the focus of Eq. 2. The fitness of an invading mutant strain *m* depends on how its susceptibility and virulence (η_mr_ and *v*_mr_) vary with the current pathogen strain *m* and the pathogen strain of the prior infection, *r* (no prior infection is represented with 0 for the strain of the prior infection, e.g., the density of hosts infected by pathogen strain *m* with no prior infection is *I_m_*_0_ with traits η*_m_*_0_ and *v*_m0_). Denoting the within-host replication rate of pathogen strain j (=*m*, *r*), as ε*_j_*, we assume that invading pathogens with higher replication rates (Eq. 3a) benefit through increased susceptibility of host populations exposed to the resident strain, i.e. ∂η_mr_/∂ε_m_ > 0 but (Eq. 3b) pay a cost in terms of higher virulence (∂*v*_mr_/∂ε_m_ > 0). We model these susceptibility and virulence relationships using the following functions, derived and parametrized from previous data (Fleming-Davies et al. 2018):

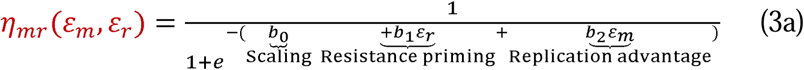

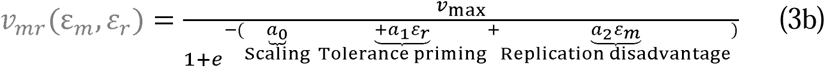

In the case of first infection of a susceptible host, the replication rate of the previous pathogen is simply 0 (ε_r_= 0 for η_m0_ or *v*_m0_). Both *b*_0_ and *a*_0_ provide baseline values for their respective functions. Both *b*_1_ and *a*_1_ represent immune priming by previous infection; *b*_1_ (resistance priming) and *a*_1_ (tolerance priming) were found to be negative, meaning that a higher replication rate of the previous infection (ε*_r_* > ε*_m_*) leads to more acquired, incomplete immunity, i.e., lower susceptibility to (η*_mr_* < η*_mm_*) and lower virulence of (*v_mr_* < *v_mm_*) reinfection (Fleming-Davies et al. 2018). Both *b*_2_ and *a*_2_ (empirically found to be positive) scale how much strongly the current strain’s replication rate (ε*_m_*) provides an advantage in terms of susceptible (*b*_2_) and a disadvantage in terms of virulence (*a*_2_).

As possible effects of provisioning, we consider provisioning that may change each of the model’s twelve parameters. In general, we assume that provisioning changes a parameter in a direction matching a direct, positive effect on the body condition of individual hosts. This directionality is based on prior empirical work showing that in the absence of disease, more feeding sites improved the body condition of captive house finch hosts as measured by antibody response or body mass change (Hawley et al. 2006). The transmission rate parameter (β), is exceptional however, as it does not necessarily reflect the body condition of the infected host.

Instead, because house finches spend more time on birdfeeders in areas with more bird feeders (Aberle et al. 2020) and bird feeder use seems linked to transmission rate (Adelman et al. 2015; Moyers et al. 2018), we assume that provisioning increases β. We consider a range of provisioning from no change in a focal parameter to ±50% change.

### Overview of model analysis

We analyze how provisioning affects pathogen virulence evolution and eco-evolutionary outcomes in the context of incomplete, acquired immunity. First, we analytically derive pathogen fitness in terms of fitness components related to infecting susceptible hosts and reinfecting recovered hosts, respectively; this derivation allows us to clearly partition pathways of how an ecological change, like provisioning, changes host traits or densities, thus changing pathogen fitness and evolution. Second, we incorporate effects of provisioning on each of the model’s parameters and solve the model numerically to explore ecological (i.e., host abundance and prevalence without pathogen evolution) and eco-evolutionary (i.e., pathogen replication rate, host abundance, and prevalence with evolution) outcomes of provisioning. We numerically find the stable resident equilibrium and analytically find invader fitness to find the eco-evolutionary equilibrium (i.e., Continuously Stable Strategy, "CSS", following Eshel 1983), toward which pathogens will evolve (see Code). We demonstrate how our decomposition of the mutant’s reproduction number allows us to understand the synergistic effects of food subsidy on different components of pathogen fitness and evolution. Further, we decompose how eco-evolutionary effects of pathogen replication rate alter prevalence and host density via immune priming vs. via the traits of the current infection. To determine the robustness of these core results, we performed a Latin hypercube search of 1000 parameter sets with each parameter within ±25% of the corresponding focal parameter value.

Third and finally, we clarify how the effects of provisioning depend on host immunity by contrasting our results with those for models without incomplete, acquired immunity or with fixed, incomplete, acquired immunity. We make the effects of provisioning comparable by setting parameters so that evolved replication rate is always the same at the level of zero provisioning for maximum comparability. For no incomplete, acquired immunity, we set *a*_1_ and *b*_1_ to 0 and adjust *a*_0_ and *b*_0_ for comparability. For fixed, incomplete, acquired immunity, we set *a*_1_ and *b*_1_ to 0 while recovered hosts have a different *a*_0_ and *b*_0_ than susceptible hosts, adjusted for comparability.

##### Box 1. Model structure and the trait functional forms of incomplete, acquired immunity

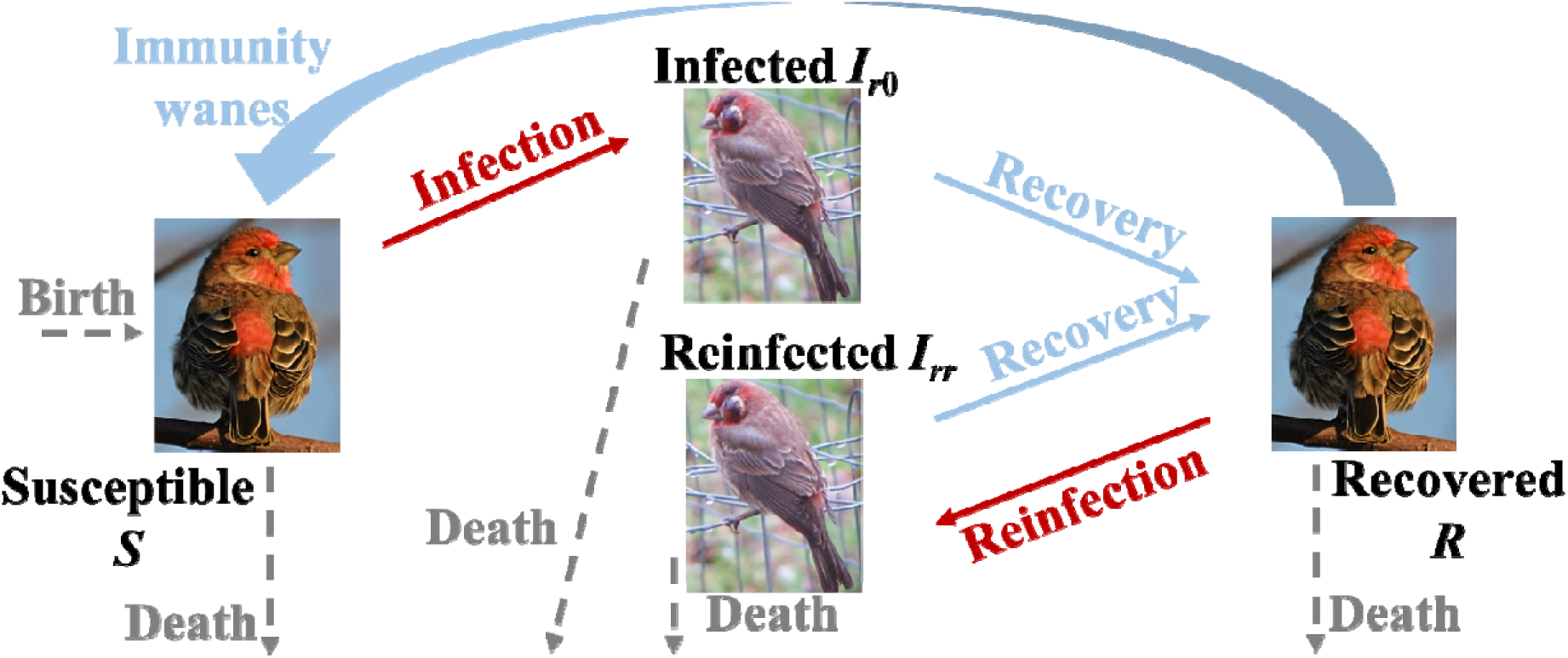

**Conceptual diagram of the disease model structure (color-coded to match Eq. 1) with a single, resident parasite strain, *r*.** Hosts are born into the susceptible class (*S*) at a fixed birth rate and die at a fixed, per-capita mortality rate. Susceptible hosts are infected by a parasite strain to become infected hosts (*I_r_*_0_); the first subscript indicates the parasite strain of the current infection, *r*, and the second indicates the parasite strain of the initial infection, which is none in this case. An infected host can recover (becoming *R*). Recovered hosts are more resistant to reinfection (which would move them into the *I_rr_*class) than susceptible hosts are to initial infection. Recovered hosts also suffer waning immunity, becoming *S* again.^1^

^1^Uninfected image: "Starry Eyed Male House Finch" by DaPuglet is licensed under CC BY-SA 2.0. To view a copy of this license, visit https://creativecommons.org/licenses/by-sa/2.0/?ref=openverse. Infected image: "Mycoplasma gallisepticum" by Ken Kneidel is marked with CC0 1.0. To view the terms, visit http://creativecommons.org/publicdomain/zero/1.0/?ref=openverse

**The functional forms of incomplete, acquired immunity for susceptibility, virulence, and their effects on the components of a mutant pathogen’s fitness (*R*_0_*^mr^*, visualized below).** We show how (A, B) susceptibility and (C, D) virulence contribute to components of fitness (E, F) across a range of mutant pathogen replication rates (ε_m_ varies along x-axis; resident replication rate kept fixed, ε*_r_* = 3.44). Each column depicts how the mutant replication rate affects parameters in hosts susceptible to the resident strain (left) or recovered from infection to the resident strain (right). Line colours depict different scenarios for the strength of how much incomplete immunity is acquired in terms of susceptibility to reinfection (parameter *b*_1_), or virulence resulting from reinfection (parameter *a*_1_): (i) ‘baseline’ (*b*_1_ = -0.41, *a*_1_ = - 0.72, black lines); (ii) resistance priming (i.e. decreased susceptibility to infection), (*b*_1_= - 0.62, orange lines); and (iii) tolerance priming (reduced virulence) (‘*a*_1_ = -1.08, blue lines). (A) Hosts are more susceptible to infection by the mutant strain when the pathogen has a higher replication rate. (B) Recovered hosts are less susceptible to reinfection than initial infection (i.e. η*_mr_* < η*_m_*_0_ for all ε*_m_*). Tolerance priming (dotted blue) does not affect susceptibility, but stronger resistance priming decreases susceptibility while simultaneously increasing its dependence on replication rate (steeper slope because the curve is less saturated). (C) Virulence increases with pathogen replication rate. (D) When recovered hosts become reinfected, they are more tolerant of infection (i.e. *v_mr_*<*v_m_*_0_ for all ε*_m_*). Stronger tolerance priming reduces virulence and reduces the sensitivity of virulence to replication rate (shallower slope) while resistance priming does not affect virulence (dashed yellow). (E) The component of the mutant pathogen’s *R*_0_ associated with susceptible hosts is maximized at low mutant replication rate (curve peaks at ε*_m_* ≈ 2). (F) The component of *R*_0_ associated with recovered hosts is maximized at much higher replication rate (ε*_m_* ≈ 4.2). Stronger tolerance priming or resistance priming both pull selection toward even higher replication rate. Gray segments are tangents at the point where invader replication rate equals resident replication rate; the segments highlight that there is no net selection given default parameter values (negative gray slope in e cancels out positive gray slope in f, implying eco-evolutionary equilibrium) but net selection for higher replication rate for either scenario with stronger priming (steeper gray slopes with stronger priming than default parameters). Here we used our focal parameter values and eco-evolutionary values for no provisioning, *S\** = 13.96, and *R\** = 11.59.

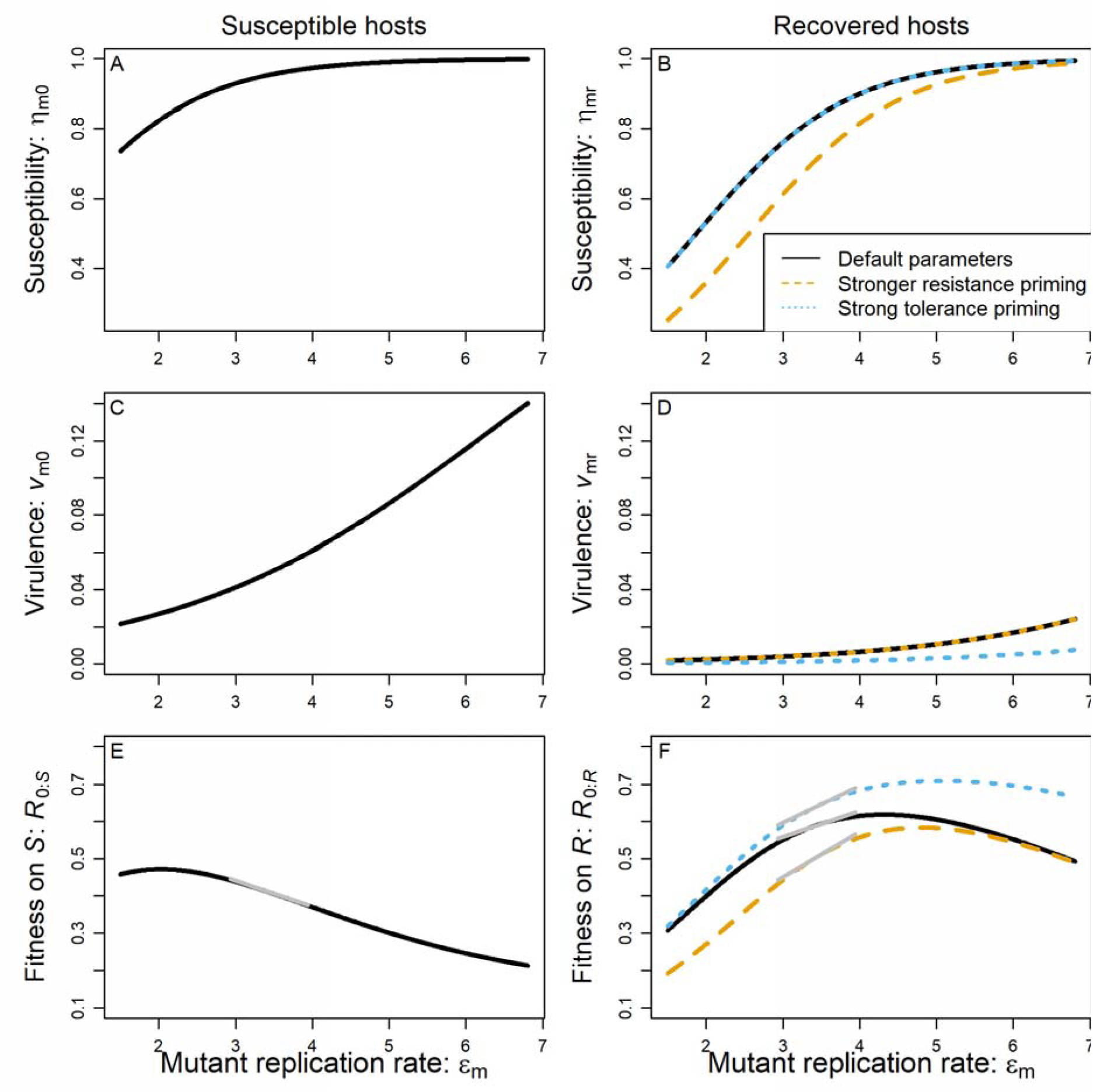

## Results

### I. Analytical derivation of pathogen fitness and partitioning pathways

We used an analytical framework to ask how reinfection dynamics will shape virulence evolution, expressing the basic reproduction number for an invading mutant in terms of the contribution from infecting hosts that are susceptible to or recovered from, infection by the resident strain (Susceptible: *R*_0:*S*_ and Recovered: *R*_0:*R*_ in Eq. 4;see Appendix for derivation):

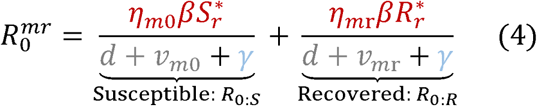

Maximizing pathogen fitness is limited by trade-offs between infections of susceptible versus recovered hosts, making it impossible to maximize both components of *R*_0_ simultaneously. The selection gradient (∂*R ^mr^*/∂ε_m_) is positive when higher replication rates (ε) lead to higher fitness and negative when lower replication rates lead to lower fitness. At eco-evolutionary equilibrium, the selection gradient must be zero, meaning the susceptible and recovered components of fitness cancel out (0 = ∂*R ^mr^*/∂ε ∂*R* /∂ε *+*∂*R* /∂ε ). Neglecting the unlikely case where they are maximized at the same value, one component of fitness must therefore favor higher replication rate (e.g., ∂*R*_0:*R*_/∂ε*_m_* > 0) while the other favors lower replication rate (∂*R*_0:*R*_/∂ε*_m_* < 0). In house finches, the traits of susceptible hosts differ from those of recovered hosts both in terms of susceptibility and virulence but also in terms of how strongly the replication rate advantage/disadvantage is realized in terms of susceptibility/virulence.

In susceptible house finches, susceptibility is generally higher (η_m0_> η_mr_) and there is a smaller advantage of pathogen replication rate than for recovered house finches (∂η_m0_/∂ε_m_ < ∂η_mr_/∂ε_m_; see height and steepness of susceptibility curves in Box 1). Both of these features cause the susceptible host component of fitness to favour lower pathogen replication rates generally, regardless of parameter values (see Appendix for proofs). Once they are infected, mortality of previously susceptible house finches is higher (*v*_m0_ > *v*_mr_) and the disadvantage of pathogen replication rate manifests more strongly than for previously recovered house finches (∂*v*_m0_/∂ε_m_ > ∂*v*_mr_/∂ε_m_; see height and steepness of virulence curves in Box 1). Higher mortality in susceptible hosts always favours higher pathogen replication rate but a stronger virulence disadvantage of replication rate always favours lower pathogen replication rate; overall, three out of four differences between susceptible and recovered hosts favour lower pathogen replication rate in susceptible house finches than recovered and thus the susceptible component of fitness peaks at lower replication rate (i.e., we find numerically ∂*R*_0:*R*_/∂ε*_m_* > 0 > ∂*R*_0:*S*_/∂ε*_m_* at ε_CSS_ for house finch-like parameter values; see gray tangents of fitness curves in Box 1). Thus, the susceptible component of fitness selects for lower pathogen replication rates while the recovered component selects for higher replication rates.

Provisioning can shift the balance of selection for pathogen replication rate through four distinct, interpretable pathways. First, provisioning often ecologically (i.e., with no evolution and thus fixed pathogen traits) decreases the equilibrium density of susceptible hosts (*S*\*) (e.g., via increased contact rate). Decreased *S*\* minimizes the weight of the *R*_0:*S*_ component of fitness (‘*S* density pathway’), so selection now favours higher replication rate. Second, provisioning often ecologically increases the density of recovered hosts (*R\**; ‘*R* density pathway’) (e.g., via slower waning of immunity), increasing the importance of the *R*_0:*R*_ component of fitness and selecting for higher replication rate. Third, provisioning can alter host ecology or immunity in ways that shift the advantage or disadvantage of replication rate within the *S* component of fitness (e.g., increased lifespan, lower δ, makes virulence disadvantages more important), altering selection for replication rate (‘*S* trade-off pathway’). Fourth and similarly, provisioning can also shift the advantage or disadvantage of replication rate in the *R* component of fitness (*R*_0:*R*_; ‘*R* trade-off pathway’). For example, provisioning that increases resistance priming (higher *b*_1_) or tolerance priming (higher *a*_1_) can shift the trade-off in the *R*_0:*R*_ component toward higher replication rates (in Box 1 dotted and dashed *R*_0:*R*_ curves peak to the right of the solid curve and typically have steeper positive slopes as shown by the gray tangents). In the following sections, we examine which pathways are most important, whether positive or negative, for provisioning-induced changes in replication rate for each model parameter.

### **II.** Ecological and eco-evolutionary effects of provisioning with incomplete, acquired immunity

Provisioning that increases host lifespan in the absence of disease (decreases δ) increases infection prevalence and selects for higher replication rate, and thus virulence. Longer lifespan ecologically, strongly decreases the density of susceptible hosts (dotted and dashed curves in Fig. 1A) while increasing the density of all other host classes (Figs. 1B-D, only weakly increased recovered host density); this effect occurs because of longer infectious periods, as fewer infected hosts die from non-disease mortality. Because increased lifespan decreases the density of susceptible hosts (Fig. 1A), it reduces the importance of the *R*_0:_ *_S_* component of fitness; the *R*_0:_ *_S_* component of fitness selects for lower replication rate so decreasing its importance favours higher replication rate (solid black increases in Fig. 1E; the *S* density pathway is strongly positive, as seen in Fig. 1F’s orange square; all other pathways are less than half its magnitude). In the ecology-only case without pathogen evolution, longer lifespan would increase the prevalence of infection and host density (dotted and dashed curves in Fig. 1A-D and Table 1). In the eco-evolutionary case with pathogen evolution of higher replication rate, prevalence and host density still increase but less so (solid curves in Fig. 1A-D and Table 1) because provisioning also selects for pathogen strains that replicate faster and are more virulent.

**Figure 1.**
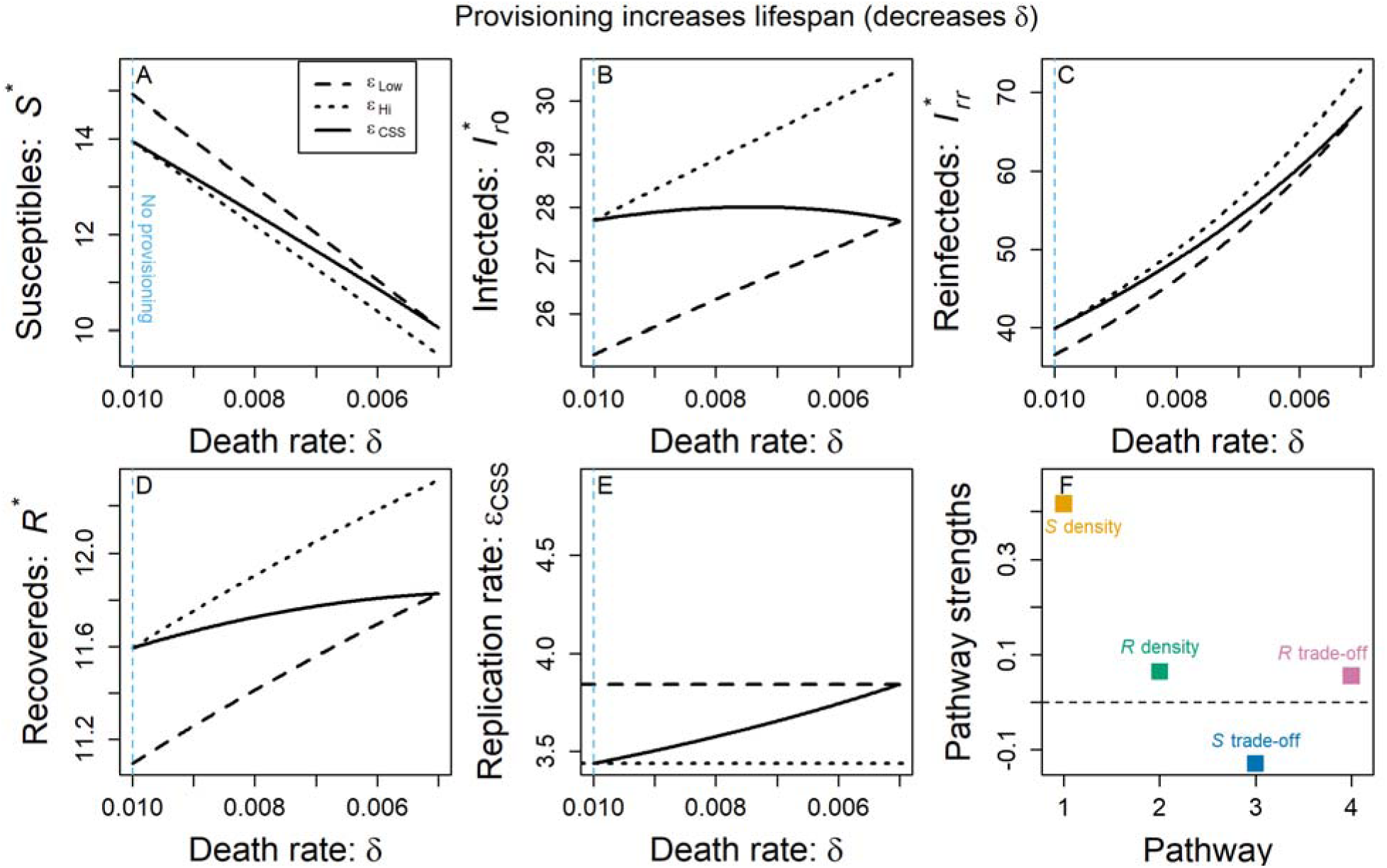
Eco-evolutionary effects of provisioning that increases host life span. We show the effect of increased lifespan and fixed, low pathogen replication rate (dotted black), fixed high replication rate (dashed black), and evolved replication rate (solid black) on (A) susceptible host density, (B) infected host density, (C) reinfected host density, (D) and recovered host density. (E) Increased lifespan selects for higher pathogen replication rate. (F) Positive pathways contribute to selection for higher pathogen replication rate while negative pathways contribute to selection for lower replication rate. The sum of pathway strengths gives the net change in ε_CSS_ seen across the provisioning range in E. Increased lifespan drives higher replication rate mainly through a positive effect via the *S* density pathway (i.e., decreased *S*\* selects for higher replication rate). The light blue, vertical dashed line shows the parameter value corresponding to default parameter values (i.e., “no provisioning”).

**Table 1.**
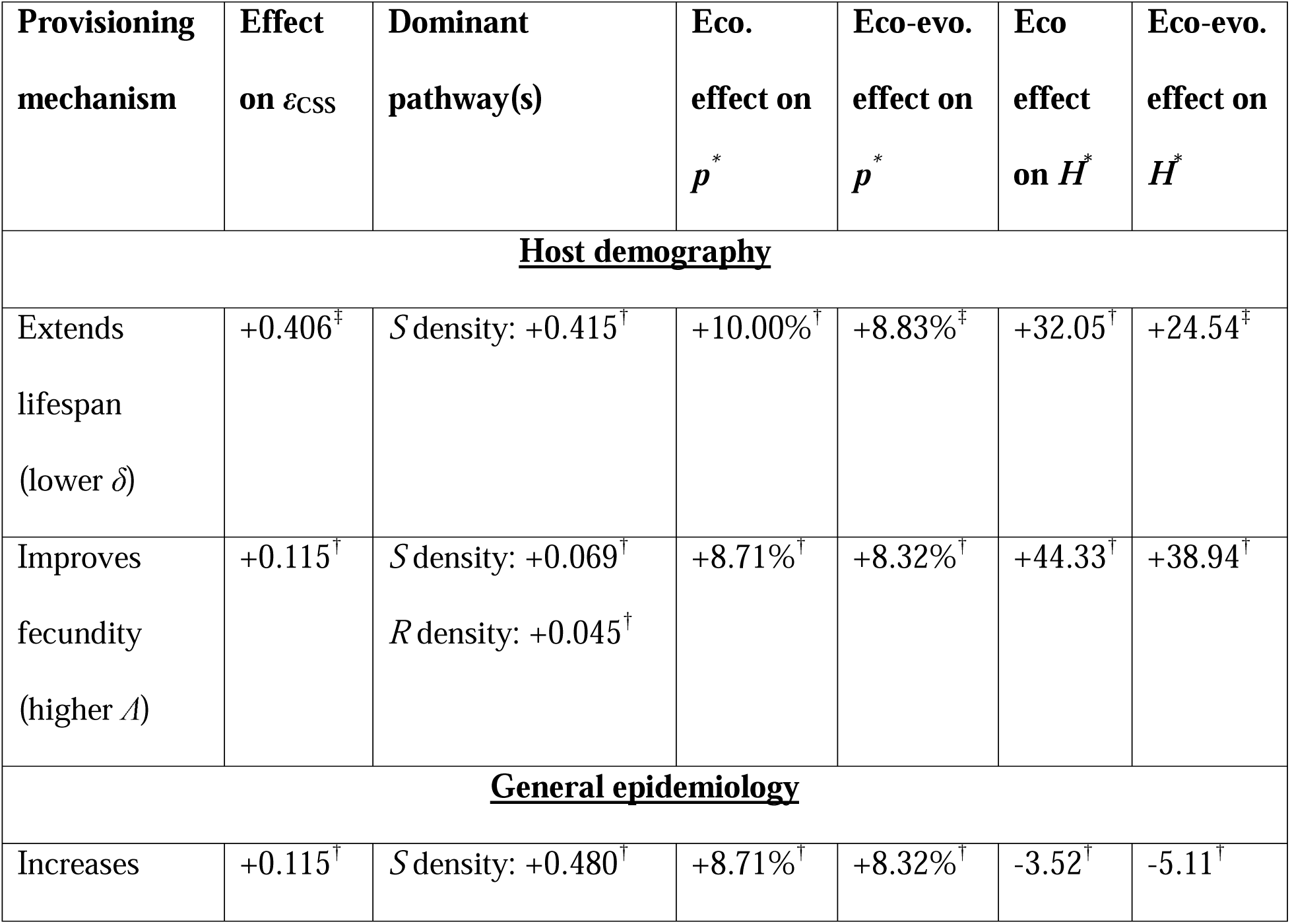
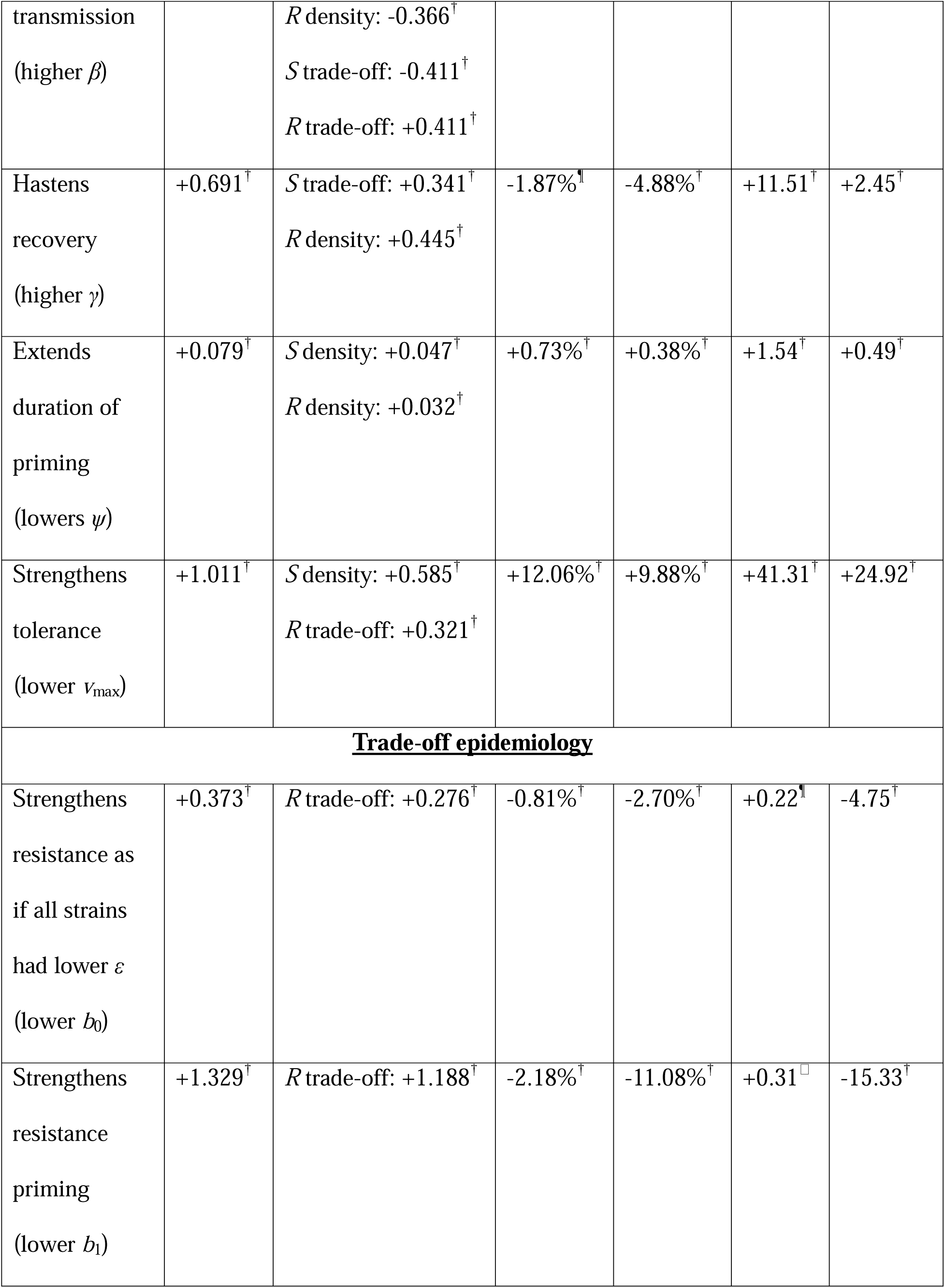
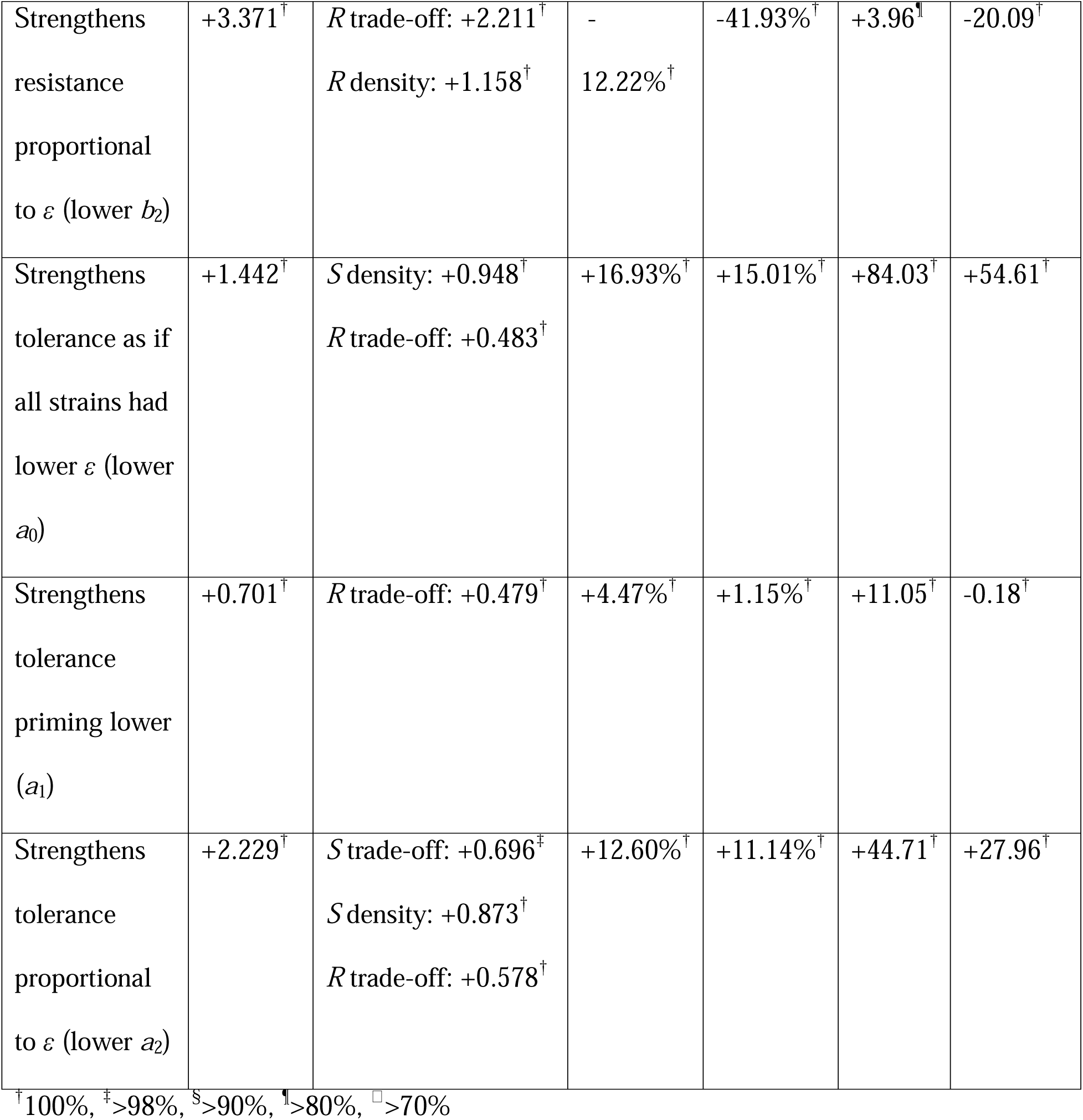
Summary of ecological and eco-evolutionary effects of different provisioning mechanisms. Provisioning acts by changing some parameter in the model, *x* (identified in the Provisioning mechanism column). We organize these mechanisms into host demography, general epidemiology, and trade-off epidemiology (i.e., non-multiplicatively alters *η*_mr_ or *v*_mr_ functions; see eq. 2). Multiple dominant pathways are shown here if their magnitude is ≥50% that of the largest magnitude pathway for that mechanism. The sign of a pathway denotes how provisioning changes the final outcome of *ε*_CSS_, e.g., provisioning that extends lifespan (decreases *δ*) decreases the density of susceptible hosts, *S*, leading to selection for higher *ε*. Ecological (no evolution) and eco-evolutionary impacts on prevalence (*p*\*) and host density (*H*\*) are also quantified. Superscripts denote how often a pattern held across the parameter search in terms of sign of effect on *ε*_CSS_, sign of the pathways that were dominant in the focal parameter set, sign of the ecological effect on *p*\*, the eco-evolutionary effect on *p*\* being more negative than the ecological effect, sign of the ecological effect on *H*\*, and the eco-evolutionary effect on *H*\* being more negative than the ecological effect (see footnote).

Provisioning may also improve host immunity, for example by strengthening how well immune priming prevents reinfection (resistance priming). Stronger resistance priming (decreased *b*_1_ in eq. 3a) ecologically reduces reinfection and thus the force of infection; these ecological effects are reflected in weakly increased susceptible host density (dotted and dashed curves in Fig. 2A), weakly increased infected host density (Fig. 2B), strongly reduced reinfected host density (Fig. 2C), and increased recovered host density (Fig. 2D). Stronger resistance priming lowers susceptibility and magnifies the susceptibility advantage of replication rate; both of these effects of resistance priming favour higher replication rate (black curve increases in Fig. 2E; see yellow curves in Box 1) for reinfecting recovered hosts (strong, positive *R* trade-off pathway as shown in Fig. 2F). In the ecology-only case without pathogen evolution, stronger resistance priming would decrease prevalence of infection and increase host density (dotted and dashed curves in Fig. 2A-D and Table 1). In the eco-evolutionary case with pathogen evolution of higher replication rate, prevalence decreases more and host density decreases instead of increasing (solid curves in Fig. 2A-D and Table 1). Thus, provisioning that increases host resistance priming can actually lead to worse outcomes for host density, due to pathogen evolution of higher replication rate and virulence.

**Figure 2.**
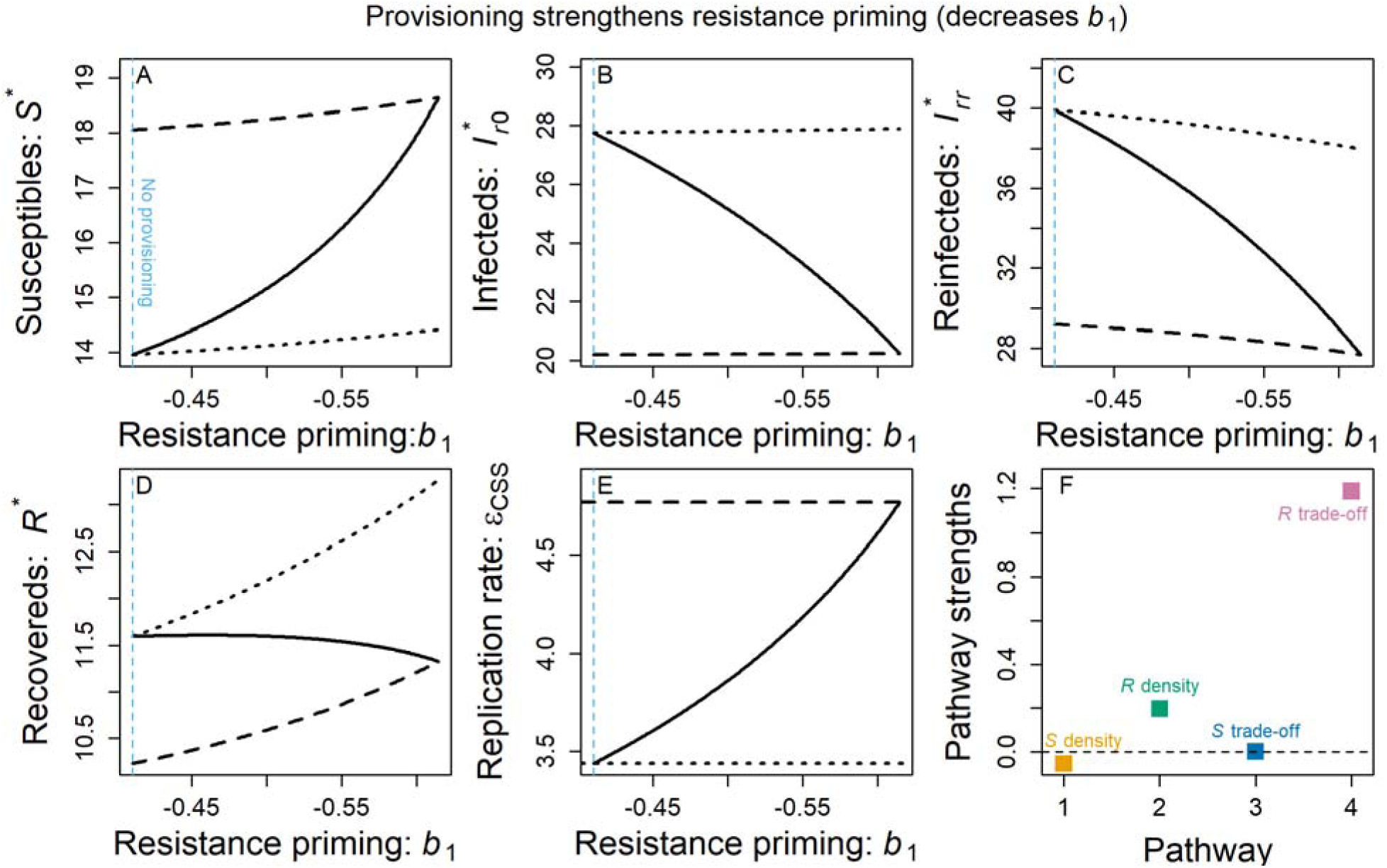
Eco-evolutionary effects of provisioning that strengthens resistance priming. We show the effect of stronger resistance priming and fixed, low pathogen replication rate (dotted black), fixed high replication rate (dashed black), and evolved replication rate (solid black) on (A) susceptible host density, (B) infected host density, (C) reinfected host density, (D) and recovered host density. (E) Stronger resistance priming strongly selects for higher pathogen replication rate. (F) Positive pathways contribute to selection for higher replication rate while negative pathways contribute to selection for lower replication rate. The sum of pathway strengths gives the net change in *ε*_CSS_ seen across the provisioning range in E. Stronger resistance priming drives higher replication rate mainly through a positive effect via the *R* trade-off pathway (i.e., stronger resistance priming makes susceptibility to reinfection lower *and* more sensitive to higher replication rate of the reinfecting strain). The light blue, vertical dashed line shows the parameter value corresponding to default parameter values (i.e., “no provisioning”).

We also considered ten other parameters that could represent provisioning mechanisms and their ecological and eco-evolutionary effects (summarized in Table 1). These include each of the different parameters of the functions controlling susceptibility and virulence (Eq. 3) as the functional form governing the interaction between host traits and pathogen traits can alter how a change in the host trait changes selection on pathogens (Williams and Day 2001). Across the entire provisioning-driven parameter range considered for each parameter, the susceptible hosts component of fitness (*R*_0:*S*_) drives selection toward lower replication rate while the recovered hosts component of fitness (*R*_0:*R*_) drives selection toward higher replication rate. Also, across all parameter ranges, provisioning selects for evolution of higher replication rate (Table 1) which, relative to the ecological effect of provisioning, causes an eco-evolutionary decrease in the prevalence of infection while also reducing host density. In some cases, these effects of pathogen evolution even create a harmful, net, eco-evolutionary effect on host density by reversing what would otherwise be, ecologically, a beneficial effect of provisioning for host density (e.g., resistance priming and tolerance priming rows of Table 1). Alternatively, there can be an eco-evolutionary decrease in infected host density (*p*\**H\**) even when this would increase without pathogen evolution (implicit in the γ row of Table 1). These results were numerically robust to our parameter search with all patterns holding in the majority of searched parameter sets and most holding in 100% of sets (see Table 1). Therefore, provisioning can drive evolution of higher pathogen virulence through many mechanisms, depressing host density.

The impact of higher pathogen replication rate on prevalence and host density is mixed because higher replication rate both protects hosts, through stronger immune priming, while simultaneously harming hosts through higher susceptibility to and virulence of a current infection. We disentangled these two conflicting effects by calculating the effect of the replication rate of the strain of the priming infection on prevalence and host density and, separately, the effect of the replication rate of the pathogen strain currently attempting to infect hosts (see attached code). Higher replication rate of the priming infection causes lower susceptibility to reinfection and lower virulence of reinfection (see Eq. 3 given *a*_1_ *<*0*, b*_1_ *<* 0, which they are empirically), leading to higher prevalence (because lower virulence matters more than lower susceptibility) and higher host density (because lower susceptibility and lower virulence both favour this). Higher replication rate of the pathogen strain currently attempting to infect hosts leads to higher susceptibility and higher virulence, leading to lower prevalence and lower host density. This pattern holds at all parameter ranges we considered when we varied provisioning; the effects of the current infection’s replication rate on prevalence and host density were usually about twice as strong as the effects of the priming infection’s replication rate. These analyses help us understand the conflicting effects of replication rate and how they lead to a net decrease in prevalence and host density (Table 1).

### **III.** Provisioning effects contrasted in different models of host immunity

The eco-evolutionary effects of provisioning depend significantly on host immunity. Without incomplete, acquired immunity (functionally a classic SIS model), provisioning can have a negative, neutral, or positive effect on evolution of replication rate (see Fig. 3A). In contrast, incomplete, acquired immunity produces a positive effect of provisioning on replication rate in every case (Fig. 3B-C). For example, provisioning that decreases mortality of infected hosts in a model without acquired immunity can drive selection for higher or lower replication rate, depending on whether decreased mortality interacts additively or multiplicatively with pathogen virulence (Williams and Day 2001) (compare δ to *v*_max_ in Fig. 3A); but with acquired immunity, the density pathways override these differences and lead all provisioning mechanisms to select for higher virulence (Fig. 3B, C). The effect of provisioning to select for higher immunity is reinforced by the reinfection biology in our focal system, and perhaps many other systems, in which variable acquired immunity depends on the replication rate of the previous infection (Fleming-Davies et al. 2018). With such variable acquired immunity, provisioning drives evolution of higher pathogen replication rate; higher replication rate, in turn, drives stronger incomplete immunity that fuels even stronger selection for higher pathogen replication rate. Thus, provisioning selects for higher replication rate most strongly with variable acquired immunity that depends on the replication rate of the previous infection (Fig. 3C). Because provisioning can select for pathogen evolution of higher replication rate, provisioning can lead to an eco-evolutionary decrease in host density, especially for variable, acquired immunity (Fig. 4).

**Figure 3.**
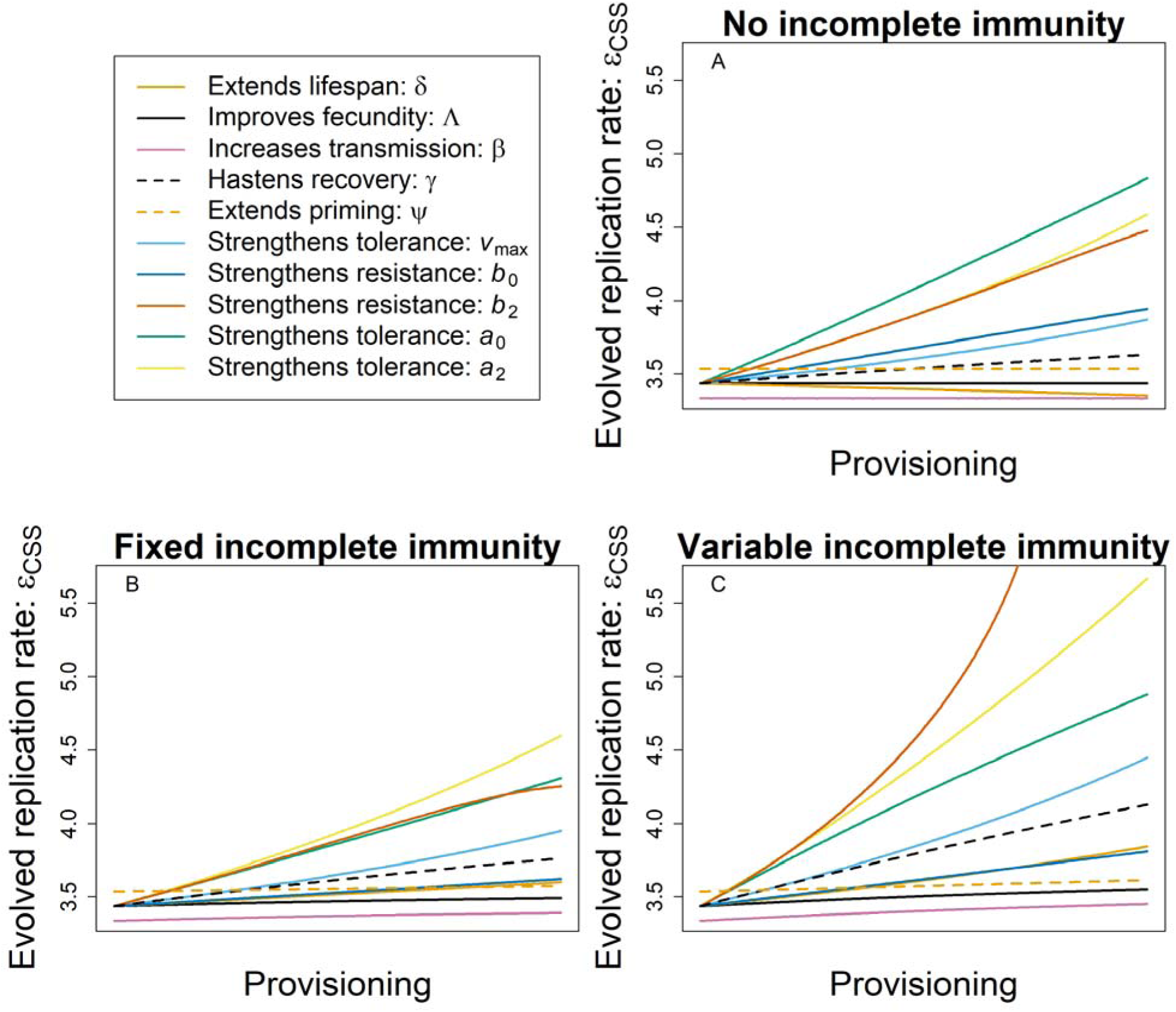
Acquired, incomplete immunity makes provisioning select for higher pathogen replication rate with negative implications for host density. Provisioning affects evolution via each parameter in each model. Provisioning increases some parameters (e.g., “Improves fecundity” means higher *Λ*) and decreases other (e.g., “Extends lifespan” means lower *δ*). (A) With no acquired, incomplete immunity, provisioning may have a positive, neutral, or negative effect on evolved replication rate, depending on which parameter responds to provisioning (positive: *γ*, *v*_max_, *b*_0_, *b*_2_, *a*_0_*, a*_2_; neutral: *Λ*, *β*, *ψ*; negative: *δ*). (B) When hosts acquire a fixed degree of incomplete immunity regardless of the pathogen strain of their initial infection (fixed so as to make Fixed and Variable incomplete immunity identical at no provisioning, see code), provisioning increases evolved replication rate for every parameter. (C) When the strength of acquired, incomplete immunity increases with the replication rate of the initial infection (Variable incomplete immunity), provisioning increases evolved replication rate for every parameter. This increase is always greater than that for Fixed incomplete immunity and almost always larger than for the corresponding case with No incomplete immunity (the only exception is that provisioning that changes *b*_0_ leads to slightly higher evolved replication rate with No incomplete immunity). The *β* curve is shifted slightly down and the *ψ* curve is shifted slightly up for visibility.

**Figure 4.**
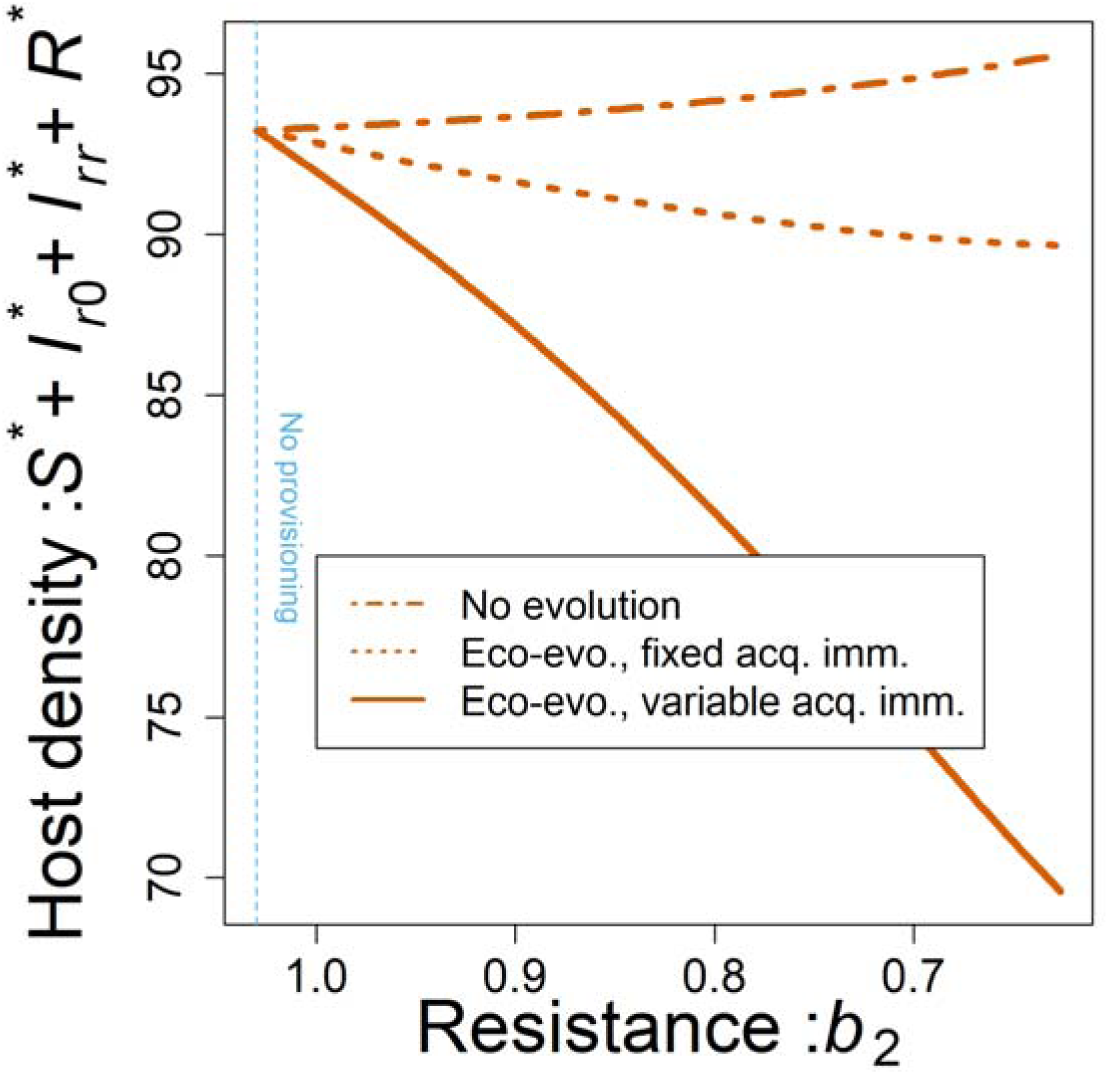
Pathogen evolution creates provisioning-driven host declines. Without evolution, provisioning typically increases host density. For example, provisioning that increases resistance (decreases *b*_2_) leads to higher equilibrium host density (dot-dashed orange). With evolution and fixed, incomplete immunity, evolution of somewhat higher pathogen replication rate causes provisioning to drive a net, eco-evolutionary decrease in host density (dotted orange). With evolution and variable, incomplete immunity, evolution of much higher replication rate causes provisioning to drive a strong, net, eco-evolutionary decrease in host density (solid orange). We do not plot densities with no incomplete immunity as we can only adjust parameters for No incomplete immunity and variable incomplete immunity to have the same evolved replication rate or host density at no provisioning (not both) and we chose to make evolved replication rates equivalent (for Fig. 3).

## Discussion

We find that anthropogenic provisioning of high-quality food may improve the body condition of wildlife yet still lead to more virulent pathogens and smaller wildlife populations, particularly in the presence of incomplete acquired immunity. Generally, provisioning that directly improves individual host body condition (as often found for high quality food: Murray et al. 2016), increases the density of recovered hosts and/or decreases the density of susceptible hosts in our model, selecting for higher virulence, selecting for higher virulence and potentially even decreasing host populations. While there is a growing expectation that incomplete immunity generally selects for higher pathogen virulence, we characterized the mixed effects of incomplete immunity in our focal system (Fleming-Davies et al. 2018). Higher tolerance in recovered hosts causes incomplete immunity to select for lower virulence. In contrast, the replication rate disadvantage of virulence manifests more weakly in recovered hosts, selecting for higher virulence. Additionally, both effects of incomplete immunity on susceptibility selected for higher virulence, leading to a net effect of incomplete immunity selecting for higher virulence.

In particular, variable, incomplete immunity that increased with the virulence of the pathogen strain of the host’s prior infection drives virulence higher than does fixed, acquired immunity; to our knowledge, this is the first direct contrast between virulence evolution in the context of variable, incomplete, acquired immunity (likely relevant to many systems: Davenport et al. 2009; Fleming-Davies et al. 2018; Moody and Downs 1955) and virulence evolution in the context of the more classic model of fixed acquired immunity (e.g., Gandon et al. 2001; Le et al. 2021). Because the effects of variable, incomplete, acquired immunity on virulence evolution was critical for understanding the outcomes from provisioning (Fig. 4), we argue there is significant value in further empirical and theoretical investigation of the multiple, possibly conflicting trait impacts of incomplete, acquired immunity on virulence evolution.

In our focal system, hosts that have recovered from infection have a substantial difference in susceptibility and other disease-related traits compared to susceptible hosts, similar to many other systems (often considered in the context of imperfect vaccination: Barnett and Civitello 2020), but we do not include other sources of heterogeneity. Previous models have found large impacts of host heterogeneity on disease spread and evolution, with heterogeneity often selecting for higher virulence (Delmas et al. 2016; Fleming-Davies et al. 2018; Gandon et al. 2001; Hoang et al. 2024; Read et al. 2015; Tate 2017). Furthermore, incomplete, acquired immunity from natural exposure or vaccination has been found to increase variation in host susceptibility in vertebrate systems including fish (Langwig et al. 2017) and the house finch disease used as the empirical system in the current study (Hawley et al. 2024). Our work emphasizes the importance of considering how these forms of host heterogeneity, such as heterogeneity within a vaccinated class, will determine the impacts of provisioning on hosts and their evolving pathogens.

Our findings advance theoretical understanding of how the impacts of provisioning depend critically on trait responses, scale, and host immunity. Previous modelling has found that provisioning may lead to ecological (no evolution) increases or decreases in pathogen prevalence, depending on which traits respond most to provisioning (Becker and Hall 2014; Erazo et al. 2022). And scale may also determine key outcomes, e.g., provisioning that improves the fitness of an individual, uninfected host (e.g., through improved fecundity) may have unfortunate consequences of raising infection prevalence at the population scale. Further, the impact of provisioning on pathogen prevalence may be positive or negative, depending on whether hosts acquire immunity after an initial infection (Becker and Hall 2014). Just as extending our consideration from the individual to population scales revealed problematic impacts of food provisioning, so too can extending from ecology-only to eco-evolutionary time scales (a conceptual extension as many pathogens evolve on ecological time scales anyway). For example, food resources might help individual hosts recover but increase infection prevalence and select for higher pathogen virulence (Hite and Cressler 2019). Previous theory of virulence evolution has focused on food resources that interact with pathogen trait trade-offs (models without acquired immunity: Hite and Cressler 2018; Hite and Cressler 2019). We bring these different theoretical threads together by considering a broad range of trait responses, individual, population, and evolutionary scales, and the impact of incomplete, acquired immunity. Further, we highlight the importance of considering host density as a key outcome. We find that provisioning that benefits individual hosts can depress host populations overall due to the interactions of pathogen evolution and acquired immunity (Fig. 4).

Our theoretical results emphasize critical areas for future empirical work. We need more laboratory measurements determining how host traits, particularly traits relevant to infection, respond to provisioning in our focal system and other systems. Further, we need measurements of prevalence, virulence, body condition, and host density across host populations receiving different levels of provisioning to help us match trait patterns to their predicted ecological or eco- evolutionary outcomes (Table 1) (see Aberle et al. 2020 for evidence of a provisioning-density association). Further, our findings put a stronger emphasis on assays that allow surveillance of the infection history of wild populations to infer the proportion of recovered hosts (such as blood antibody titers). In addition to field work, laboratory epidemics can assay the eco-evolutionary impacts of resource availability (Lopez-Pascua et al. 2014; Walsman et al. 2023); flock experiments with captive house finches (Moyers et al. 2018) could be extended with simulated mortality and competition among pathogen strains to rigorously test theoretical predictions. Such empirical work will strengthen our understanding of how provisioning may reshape disease outcomes for host populations.

Overall, our results add to the growing literature raising concerns for how anthropogenic change will alter disease outcomes. There are many anthropogenic effects on diseases of wildlife, which can stimulate pathogen responses that are hard to predict and manage due to the speed of pathogen spread and evolution (Rogalski et al. 2017). Pathogen evolution, in turn, may rapidly impact host population dynamics or even lead to host jumps, in extreme cases (Tan et al. 2024). Such dynamic pathogen responses to anthropogenic change can easily lead to harmful or unanticipated consequences for wildlife, even if anthropogenic provisioning strengthens host immunity. We need to anticipate these outcomes, e.g., if we wish to reduce infected host density to prevent disease spill-over or maintain total host density to conserve a species. This need is exemplified by provisioning that directly improves host body condition but can indirectly lead to deadlier diseases and smaller wildlife populations.

## Appendix

### Parameter values and meanings

**Table A1.** Parameter values and meanings. For each parameter (symbol in first column), we provide a verbal description of the meaning of the parameter (second column), and the values and that correspond to our no provisioning scenario. All parameter values follow (Fleming-Davies et al. 2018).

| Parameter | Meaning | Value |
| --- | --- | --- |
| $\Lambda$ | Host fecundity | 2.5 |
| $b_0$ | Factor controlling susceptibility | -0.52 |
| $b_1$ | Sensitivity of resistance priming to replication rate | -0.41 |
| $b_2$ | Sensitivity of susceptibility to replication rate | 1.03 |
| $\beta$ | Transmission rate | 0.00275 |
| $a_0$ | Factor controlling virulence | -3.09 |
| $a_1$ | Sensitivity of tolerance priming to replication rate | -0.72 |
| $a_2$ | Sensitivity of virulence to replication rate | 0.49 |
| $v_{\max}$ | Maximum virulence | 0.25 |
| $\varepsilon$ | Replication rate of pathogens | Evolves and $\varepsilon_{\text{CSS}} = 3.43$<br>without provisioning |
| $\delta$ | Per-capita death rate of hosts without disease | 0.01 |
| $\psi$ | Rate of immunity waning | 0.01 |
| $\gamma$ | Rate of recovery from infection | 0.03 |

#### Proof that *R ^mr^* can be expressed as *R + R* components (Eq. 4)

We calculate the fitness of an invading pathogen strain as *R*_0_ using the Next Generation Matrix approach (Hurford et al. 2009). We construct this matrix using the equations for a rare mutant *m*’s d*I_m_*_0_/d*t*, d*I_m_*_r_/d*t*, and d*I_m_*_m_/d*t* (see Eq. 2). The Next Generation Matrix takes the form shown in Eq. A1a so that the eigenvalues are elements of the vector shown in Eq. A1b, proving the expression in Eq. 4.

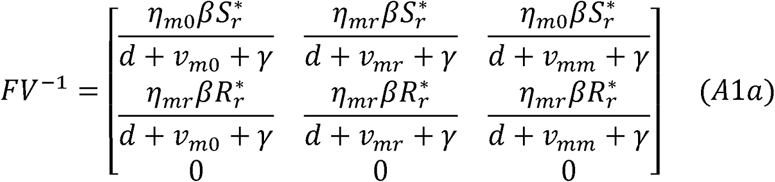

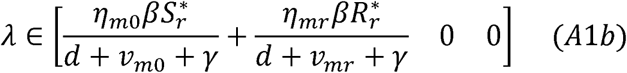

### Proof of how acquired immunity alters selection

Consider a population at eco-evolutionary equilibrium (*ε*_CSS: no acq. imm._, implying eq. A2a) in which recovered hosts possess no acquired immunity (η_m0_ = η_mr_, *v*_m0_ = *v*_mr_, implying eq. A2b). We can derive that the recovered component of fitness is also zero at equilibrium (i.e., eqs. A2a and A2b imply A2c).

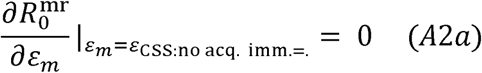

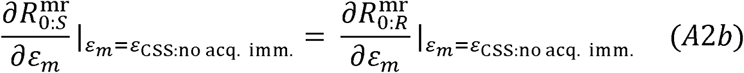

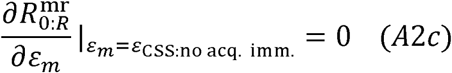

We can inspect how some acquired immunity, instead of none, would affect the impact of replication rate on the recovered component of fitness.

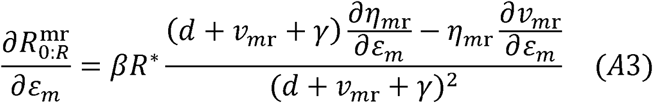

In the susceptibility-virulence trade-off, we assumed that both of the partial derivatives on the right-hand side of eq. A3 were positive. Consider a change in the system such that there is now some acquired immunity, e.g., η_m0_ remains the same but η_mr_ decreases by a small amount. We can easily see from eq. A3 that a decrease in η_mr_, as occurs in house finches with acquired immunity, increases ∂*R*_0:*R*_^mr^/∂*ε_m_* |*ε_m=_ε*_CSS: no acq. imm._ from 0 to positive while ∂*R*_0:*S*_^mr^/∂*ε_m_* |*ε_m=_ε*_CSS: no acq. imm._ Is still 0. If we assume a continuous change in *ε*_CSS_as η_mr_ decreases from the no acquired immunity case to a slightly smaller value in the acquired immunity case, then clearly *ε*_CSS: no acq. imm_< *ε*_CSS: acq. imm_; in other words, a decrease in the susceptibility of recovered hosts compared to susceptible hosts (due to acquired immunity) selects for higher replication rate. The same reasoning can be applied to determine whether the other empirically characterized aspects of acquired immunity in house finches contribute to selection for higher or lower replication rate; higher ∂η^mr^/∂*ε_m_* selects for higher replication rate, lower *v*^mr^ selects for lower replication rate, and lower ∂*v*^mr^/∂*ε_m_* selects for higher replication rate.

### Further mathematical details on how the four pathways are calculated

We focus on how the behaviour of the selection gradient (*G* = ∂*R*_0_/∂*ε*_m_|*ε*_m=_*ε*_r_), captures how a change in replication rate will alter fitness. An evolutionarily stable point (known as a “Continuously Stable Strategy” or “CSS”) is a replication rate, *ε*_CSS_, at which evolution will stop (*G* = 0, evolutionary singular point), a local fitness optimum (∂/∂*ε*_m_ *G* < 0, evolutionary stability), and toward which a population will evolve (d/d*ε G* < 0, convergence stability). When provisioning changes some parameter *x*, provisioning changes *ε*_CSS_ by changing the selection gradient (eq. A4a). We can partition the ways by which provisioning can change *G* into four pathways with clear interpretation (eq. A4b):

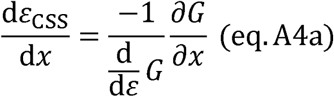

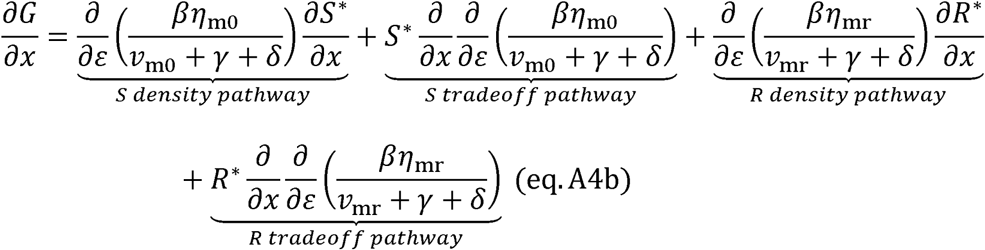

Considering susceptible hosts, provisioning may either change selection by changing their density [*S* density pathway] or by altering the balance of susceptibility and virulence when infecting susceptible hosts [*S* tradeoff pathway]. Similarly for recovered hosts, provisioning may either change selection by changing their density [*R* density pathway] or by altering the balance of susceptibility and virulence when infecting recovered hosts [*R* tradeoff pathway]. These pathways help us interpret the overall effects of provisioning on pathogen evolution.

### Detailed eco-evolutionary results for other provisioning effects (Figs. A1-A10)

**Figure A1.**
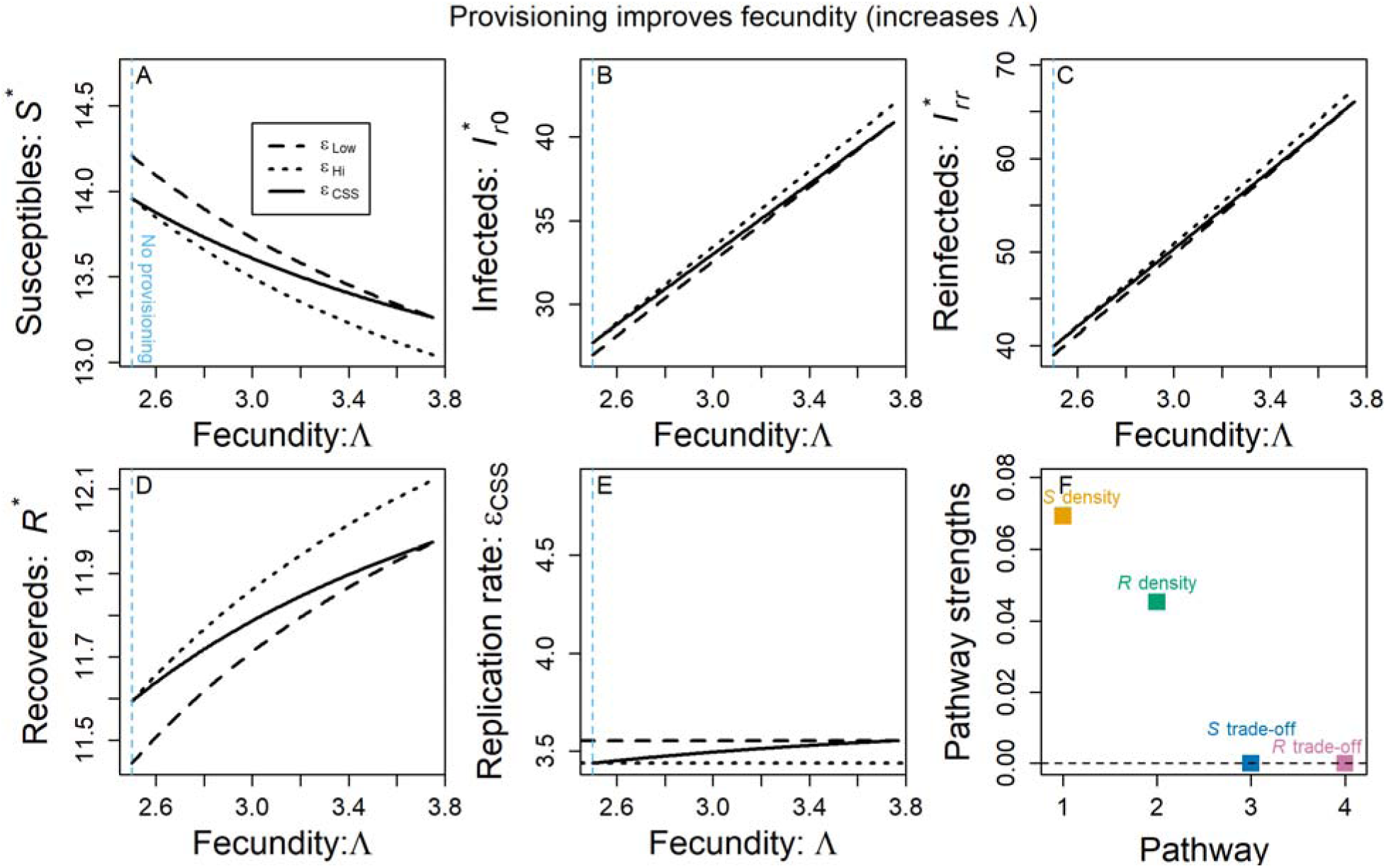
Eco-evolutionary effects of provisioning that improves fecundity. We show the effect of higher fecundity and fixed, low replication rate (dotted black), fixed high replication rate (dashed black), and evolved replication rate (solid black) on (A) susceptible host density, (B) infected host density, (C) reinfected host density, (D) and recovered host density. (E) Higher fecundity weakly selects for higher pathogen replication rate. (F) Positive pathways contribute to selection for higher replication rate while negative pathways contribute to selection for lower replication rate. The sum of pathway strengths gives the net change in *ε*_CSS_ seen across the provisioning range in Fig. A1E. Higher fecundity drives higher replication rate only through positive effects via the *S* density and *R* density pathways (i.e., higher fecundity decreases *S*\* and increases *R*\*, both of which select for higher *ε*). The light blue, vertical dashed line shows the parameter value corresponding to default parameter values (i.e., “no provisioning”).

**Figure A2.**
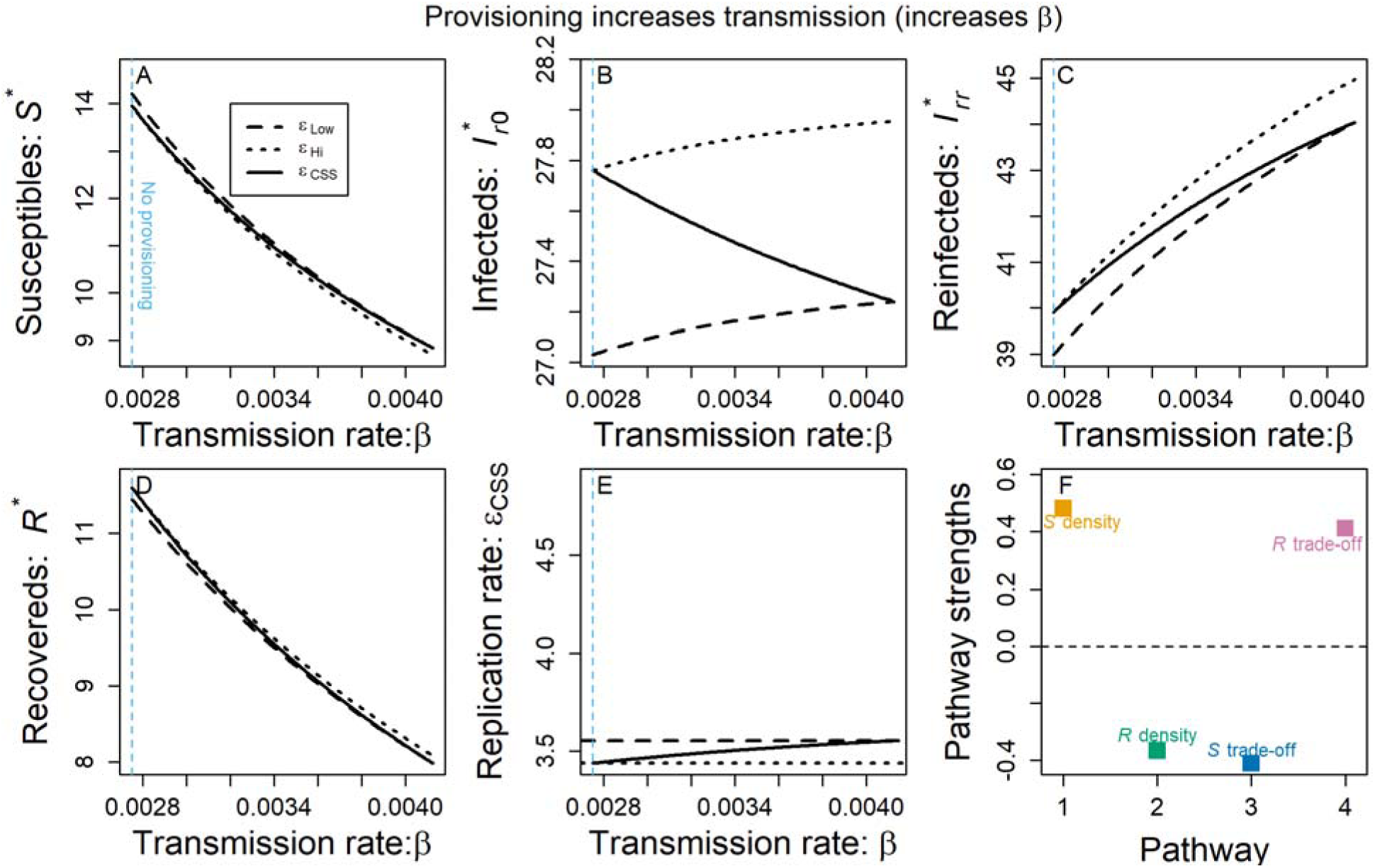
Eco-evolutionary effects of provisioning that increases transmission. We show the effect of higher transmission rate and fixed, low replication rate (dotted black), fixed high replication rate (dashed black), and evolved replication rate (solid black) on (A) susceptible host density, (B) infected host density, (C) reinfected host density, (D) and recovered host density. (E) Higher transmission rate weakly selects for higher pathogen replication rate. (F) Positive pathways contribute to selection for higher replication rate while negative pathways contribute to selection for lower replication rate. The sum of pathway strengths gives the net change in *ε*_CSS_ seen across the provisioning range in Fig. A2E. Higher transmission rate drives higher replication rate through positive effects via the *S* density and *R*-tradeoff pathways. The light blue, vertical dashed line shows the parameter value corresponding to default parameter values (i.e., “no provisioning”).

**Figure A3.**
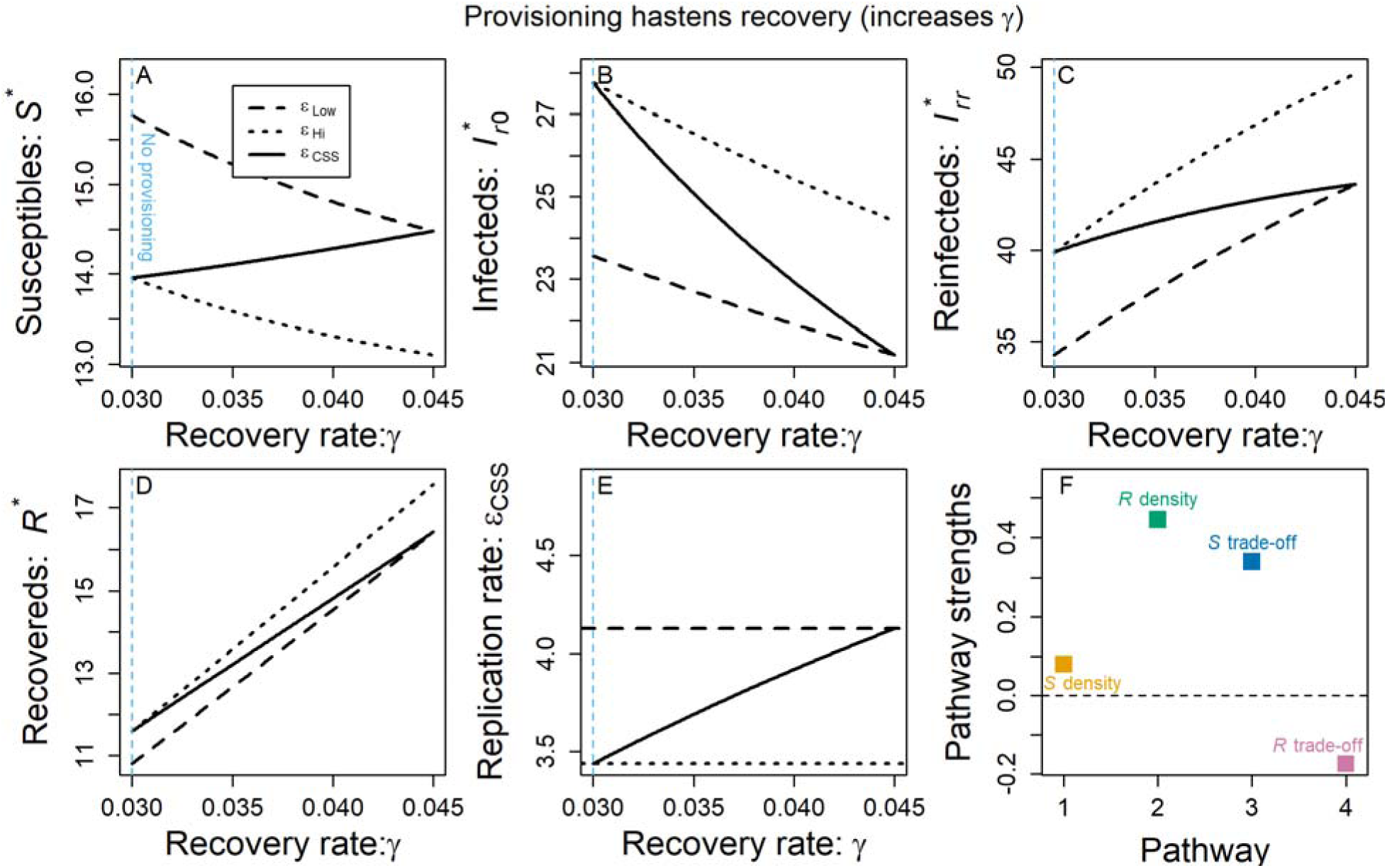
**Eco-evolutionary effects of provisioning that increases recovery. Γ**

**Figure A4.**
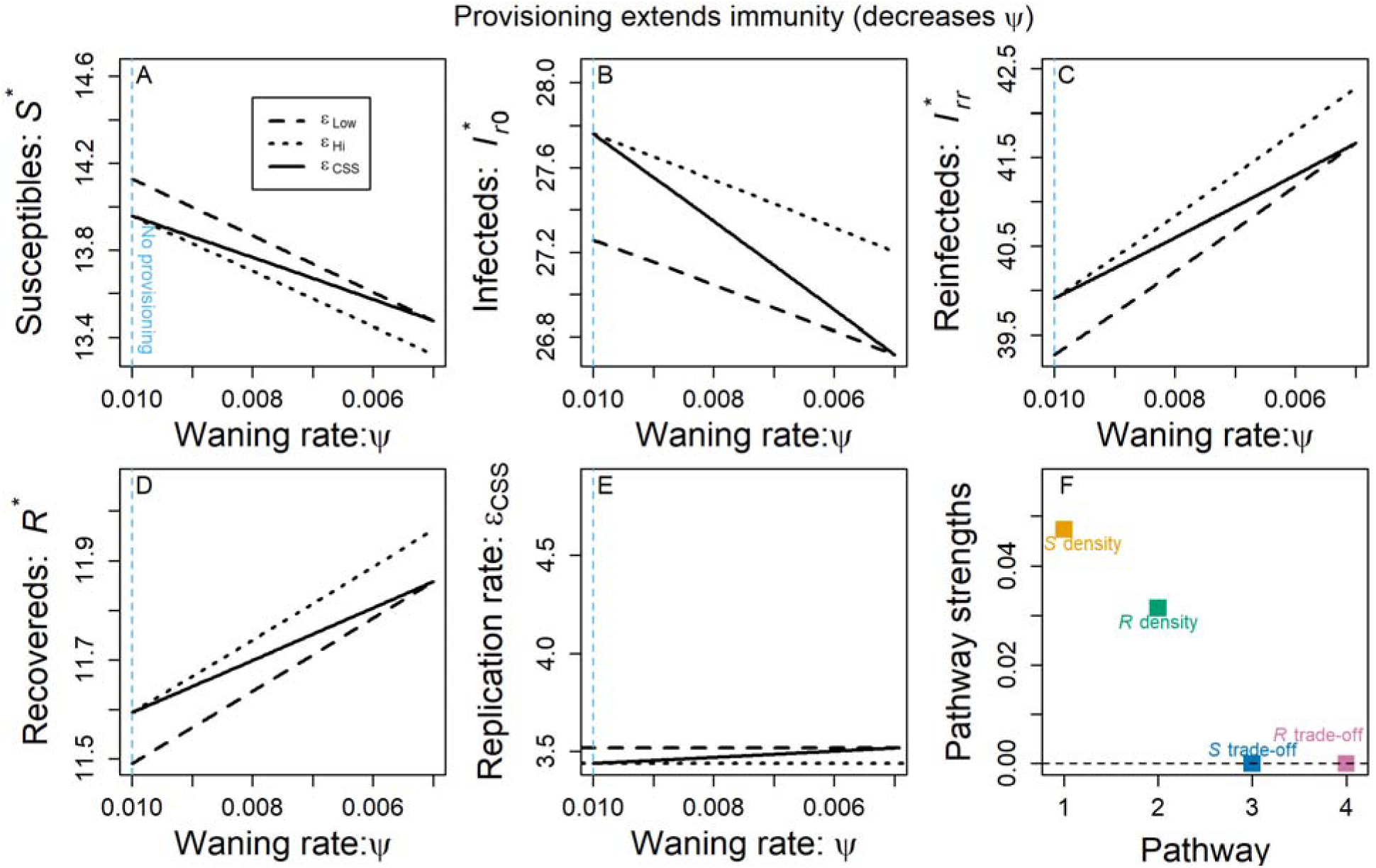
Eco-evolutionary effects of provisioning that extends immune priming. We show the effect of extended priming and fixed, low replication rate (dotted black), fixed high replication rate (dashed black), and evolved replication rate (solid black) on (A) susceptible host density, (B) infected host density, (C) reinfected host density, (D) and recovered host density. (E) Extended priming weakly selects for higher pathogen replication rate. (F) Positive pathways contribute to selection for higher replication rate while negative pathways contribute to selection for lower replication rate. The sum of pathway strengths gives the net change in *ε*_CSS_ seen across the provisioning range in Fig. A4E. Extended priming drives higher replication rate only through positive effects via the *S* density and *R* density pathways (i.e., extended priming decreases *S*\* and increases *R*\*, both of which select for higher *ε*). The light blue, vertical dashed line shows the parameter value corresponding to default parameter values (i.e., “no provisioning”).

**Figure A5.**
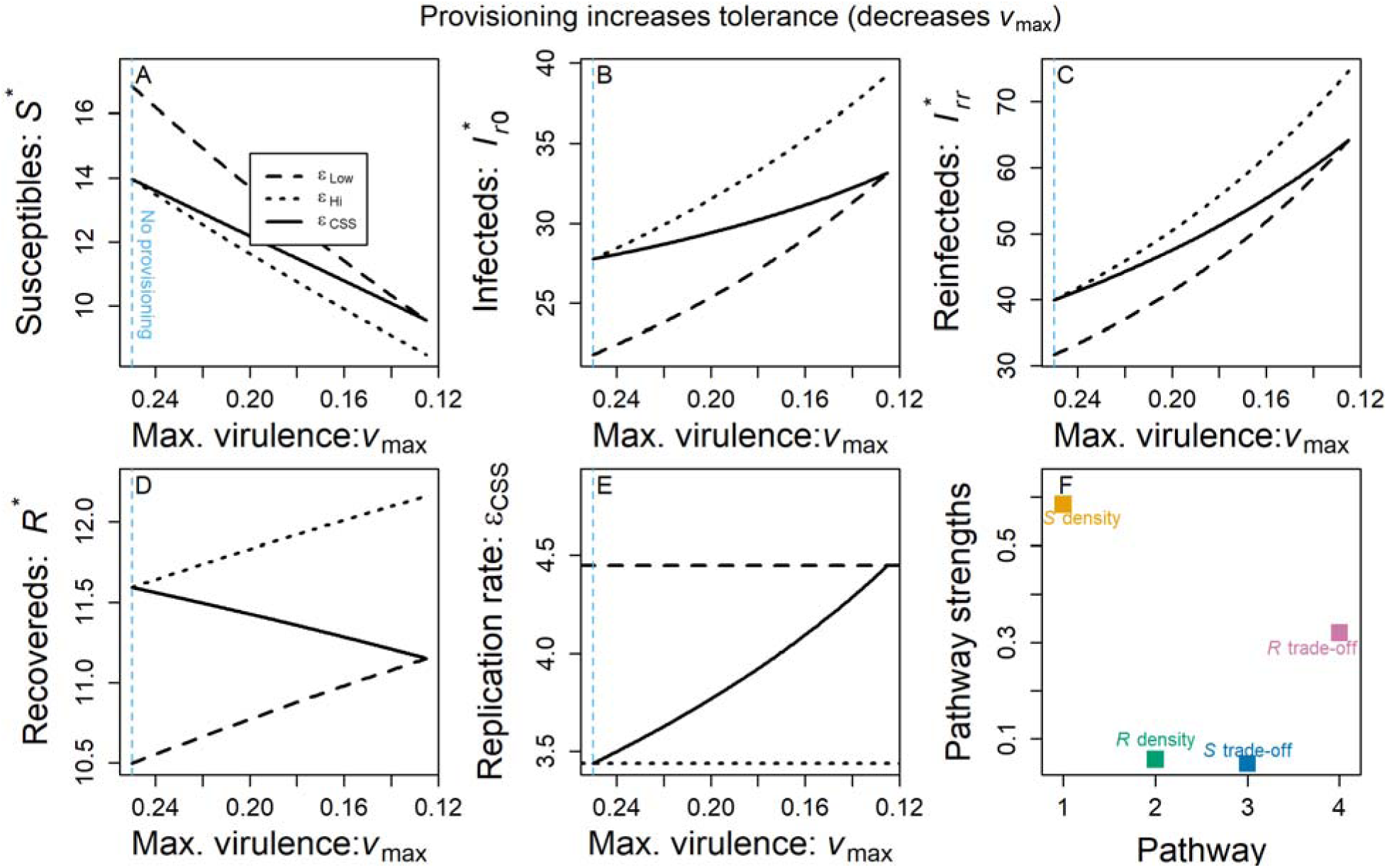
Eco-evolutionary effects of provisioning that strengthens tolerance. We show the effect of higher tolerance and fixed, low replication rate (dotted black), fixed high replication rate (dashed black), and evolved replication rate (solid black) on (A) susceptible host density, (B) infected host density, (C) reinfected host density, (D) and recovered host density. (E) Higher tolerance selects for higher pathogen replication rate. (F) Positive pathways contribute to selection for higher replication rate while negative pathways contribute to selection for lower replication rate. The sum of pathway strengths gives the net change in *ε*_CSS_ seen across the provisioning range in Fig. A5E. Higher tolerance drives higher replication mainly through positive effects via the *S* density and *R* trade-off pathways. The light blue, vertical dashed line shows the parameter value corresponding to default parameter values (i.e., “no provisioning”).

**Figure A6.**
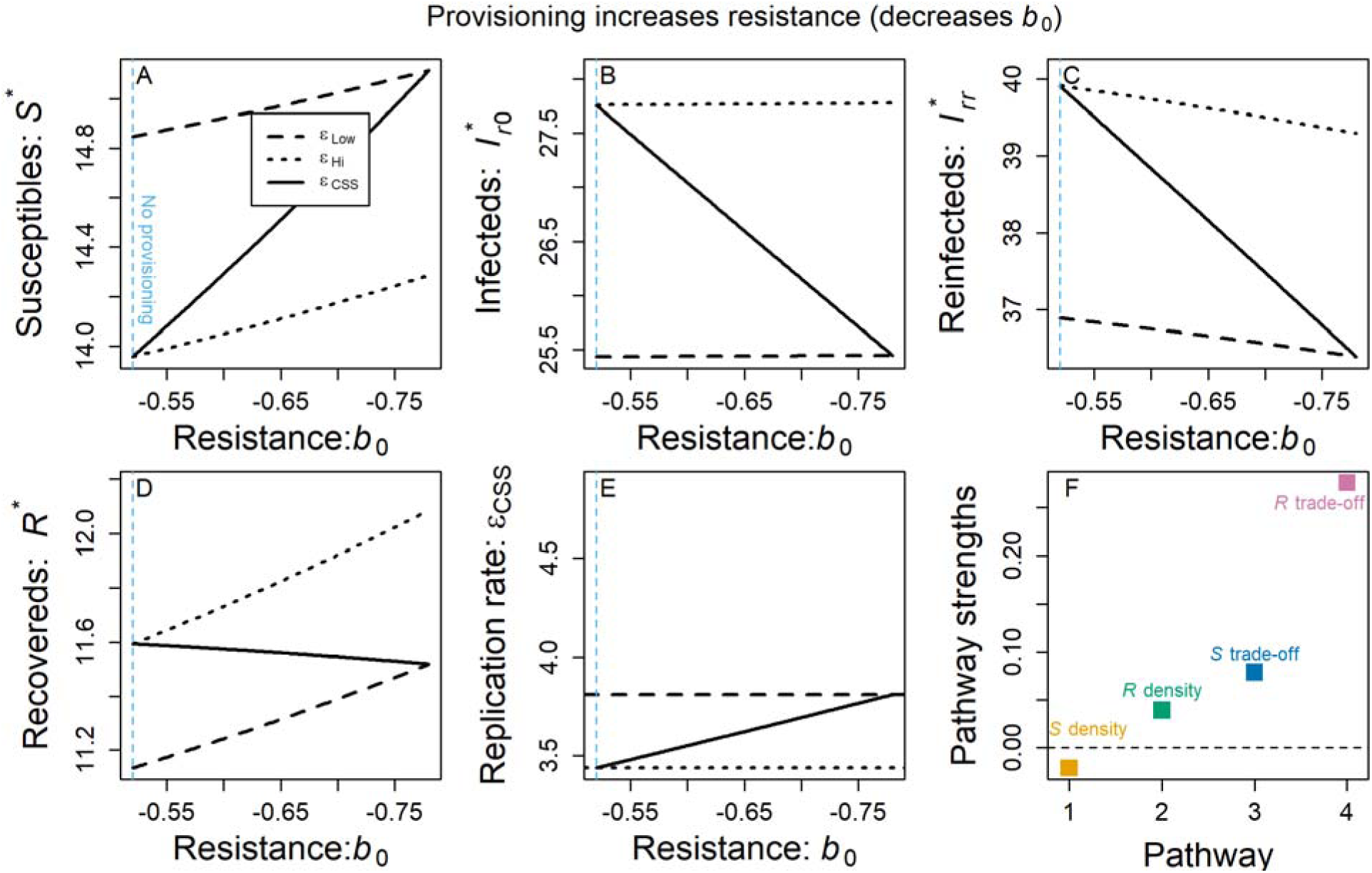
Eco-evolutionary effects of provisioning that strengthens resistance. We show the effect of stronger resistance (through lower *b*_0_) and fixed, low replication rate (dotted black), fixed high replication rate (dashed black), and evolved replication rate (solid black) on (A) susceptible host density, (B) infected host density, (C) reinfected host density, (D) and recovered host density. (E) Stronger resistance weakly selects for higher pathogen replication rate. (F) Positive pathways contribute to selection for higher replication rate while negative pathways contribute to selection for lower replication rate. The sum of pathway strengths gives the net change in *ε*_CSS_ seen across the provisioning range in Fig. A6E. Stronger resistance drives higher replication rate mainly through a positive *R* trade-off pathway. The light blue, vertical dashed line shows the parameter value corresponding to default parameter values (i.e., “no provisioning”).

**Figure A7.**
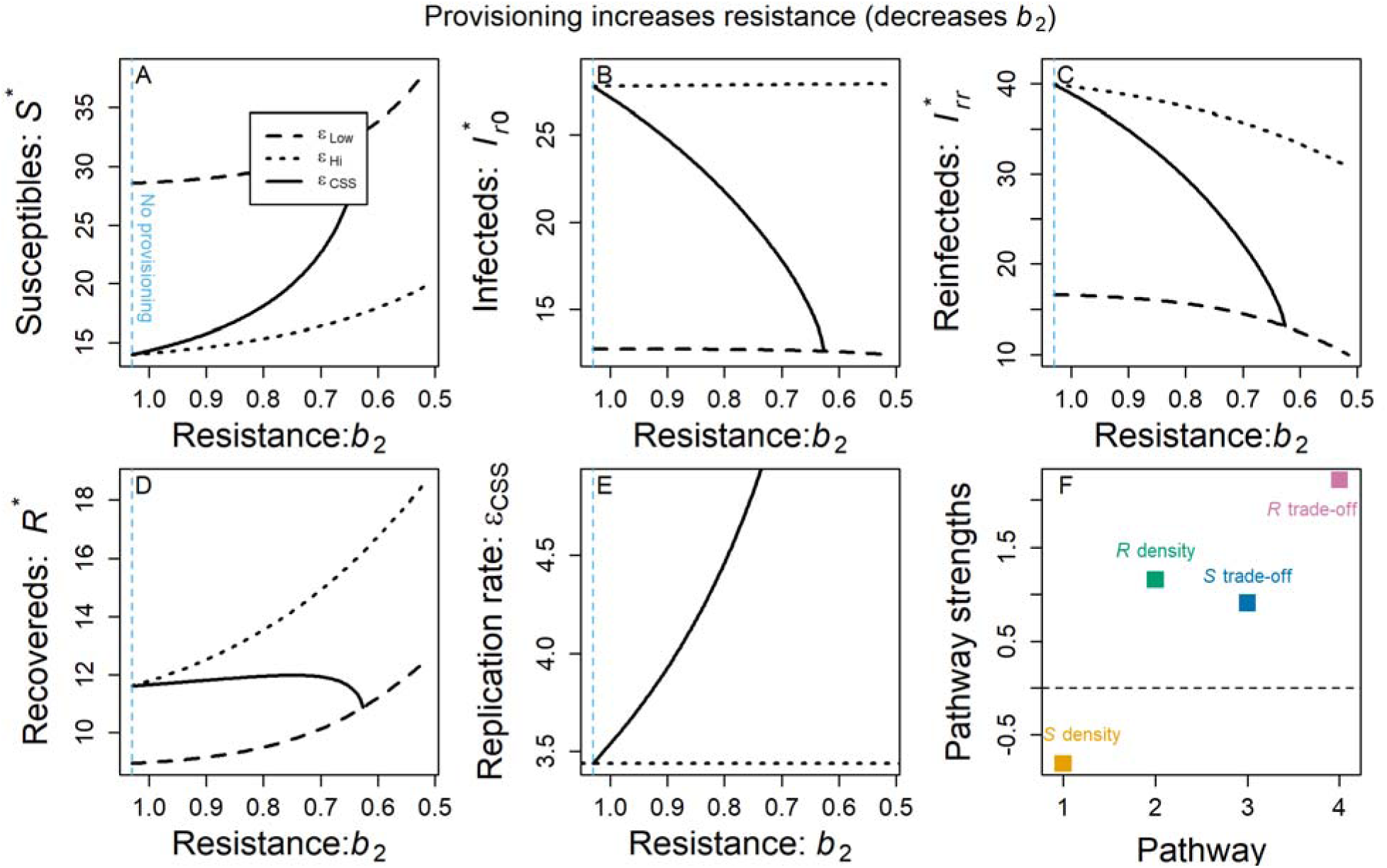
Eco-evolutionary effects of provisioning that strengthens resistance. We show the effect of stronger resistance (through lower *b*_2_) and fixed, low replication rate (dotted black), fixed high replication rate (dashed black), and evolved replication rate (solid black) on (A) susceptible host density, (B) infected host density, (C) reinfected host density, (D) and recovered host density. (E) Stronger resistance strongly selects for higher pathogen replication rate. (F) Positive pathways contribute to selection for higher replication rate while negative pathways contribute to selection for lower replication rate. The sum of pathway strengths gives the net change in *ε*_CSS_ seen across the provisioning range in Fig. A7E. Stronger resistance drives higher replication rate through positive *R* density, *S* trade-off, and *R* trade-off pathways. The light blue, vertical dashed line shows the parameter value corresponding to default parameter values (i.e., “no provisioning”).

**Figure A8.**
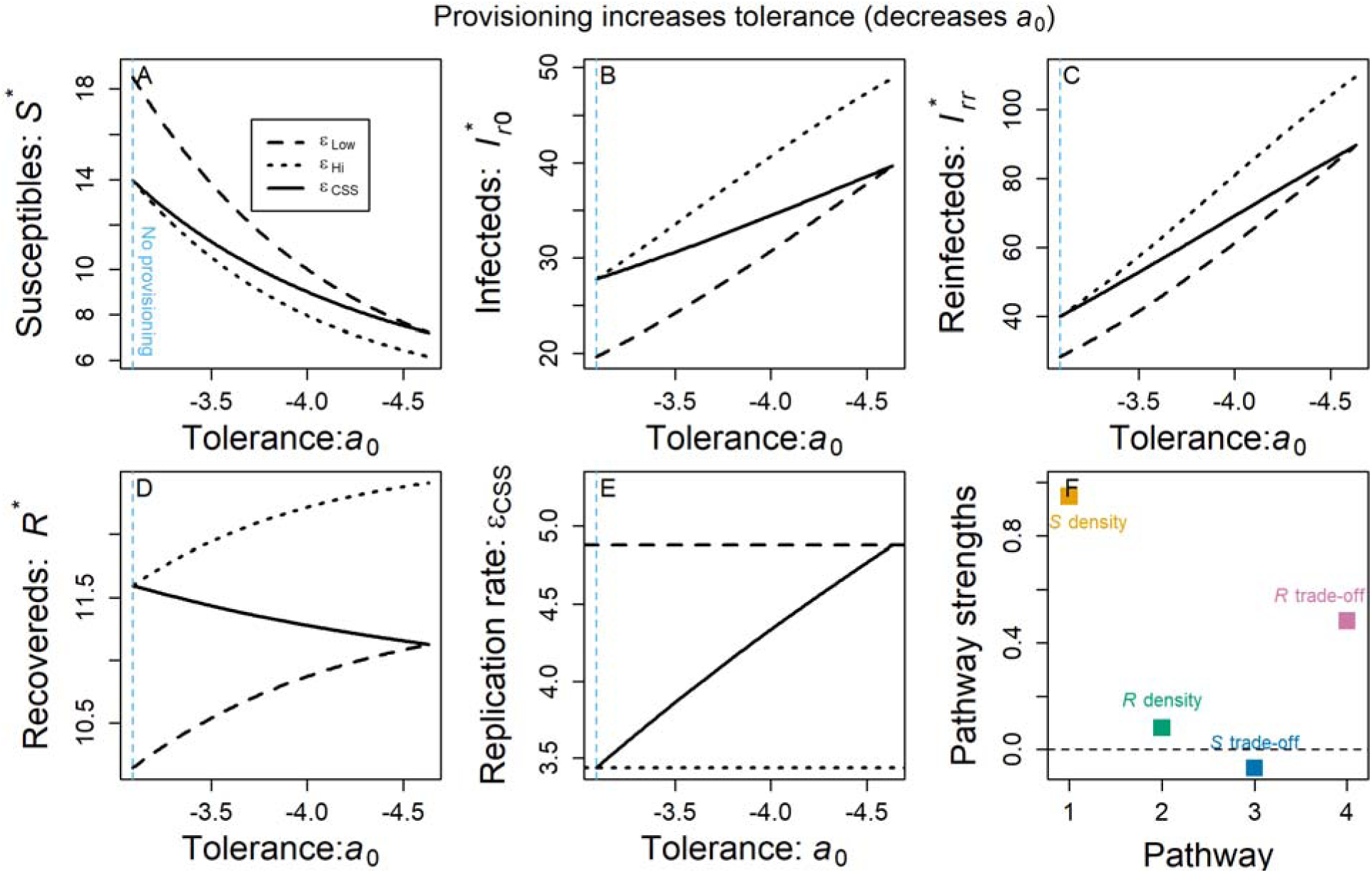
Eco-evolutionary effects of provisioning that strengthens tolerance. We show the effect of stronger tolerance (through lower *a*_0_) and fixed, low replication rate (dotted black), fixed high replication rate (dashed black), and evolved replication rate (solid black) on (A) susceptible host density, (B) infected host density, (C) reinfected host density, (D) and recovered host density. (E) Stronger tolerance strongly selects for higher pathogen replication rate. (F) Positive pathways contribute to selection for higher replication rate while negative pathways contribute to selection for lower replication rate. The sum of pathway strengths gives the net change in *ε*_CSS_ seen across the provisioning range in Fig. A8E. Stronger tolerance drives higher replication rate mainly through positive *S* density and *R* trade-off pathways. The light blue, vertical dashed line shows the parameter value corresponding to default parameter values (i.e., “no provisioning”).

**Figure A9.**
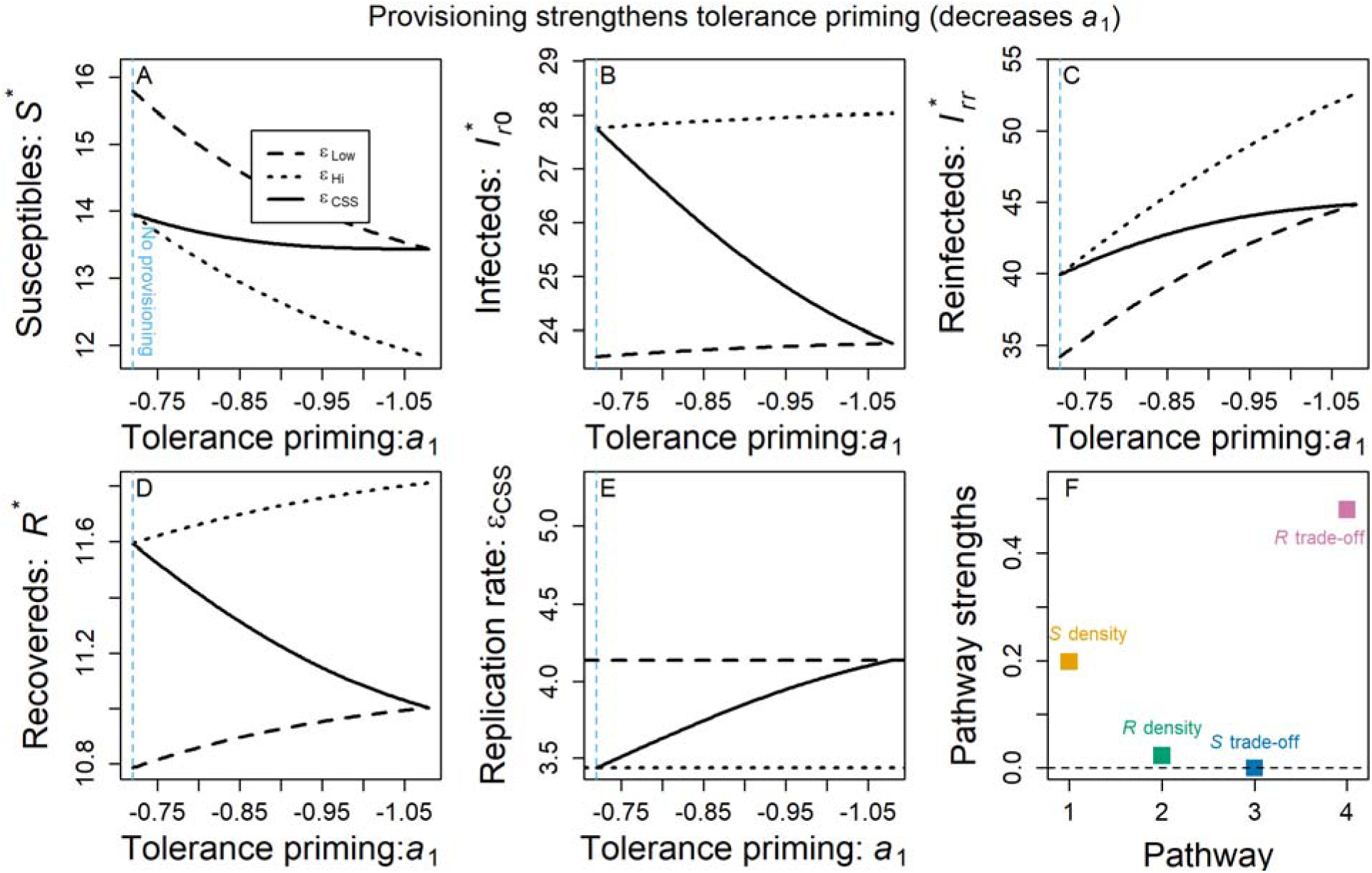
Eco-evolutionary effects of provisioning that strengthens tolerance priming. We show the effect of stronger tolerance priming (through lower *a*_1_) and fixed, low replication rate (dotted black), fixed high replication rate (dashed black), and evolved replication rate (solid black) on (A) susceptible host density, (B) infected host density, (C) reinfected host density, (D) and recovered host density. (E) Stronger tolerance priming selects for higher pathogen replication rate. (F) Positive pathways contribute to selection for higher replication rate while negative pathways contribute to selection for lower replication rate. The sum of pathway strengths gives the net change in *ε*_CSS_ seen across the provisioning range in Fig. A9E. Stronger tolerance priming drives higher replication rate mainly through positive *S* density and *R* trade-off pathways. The light blue, vertical dashed line shows the parameter value corresponding to default parameter values (i.e., “no provisioning”).

**Figure A10.**
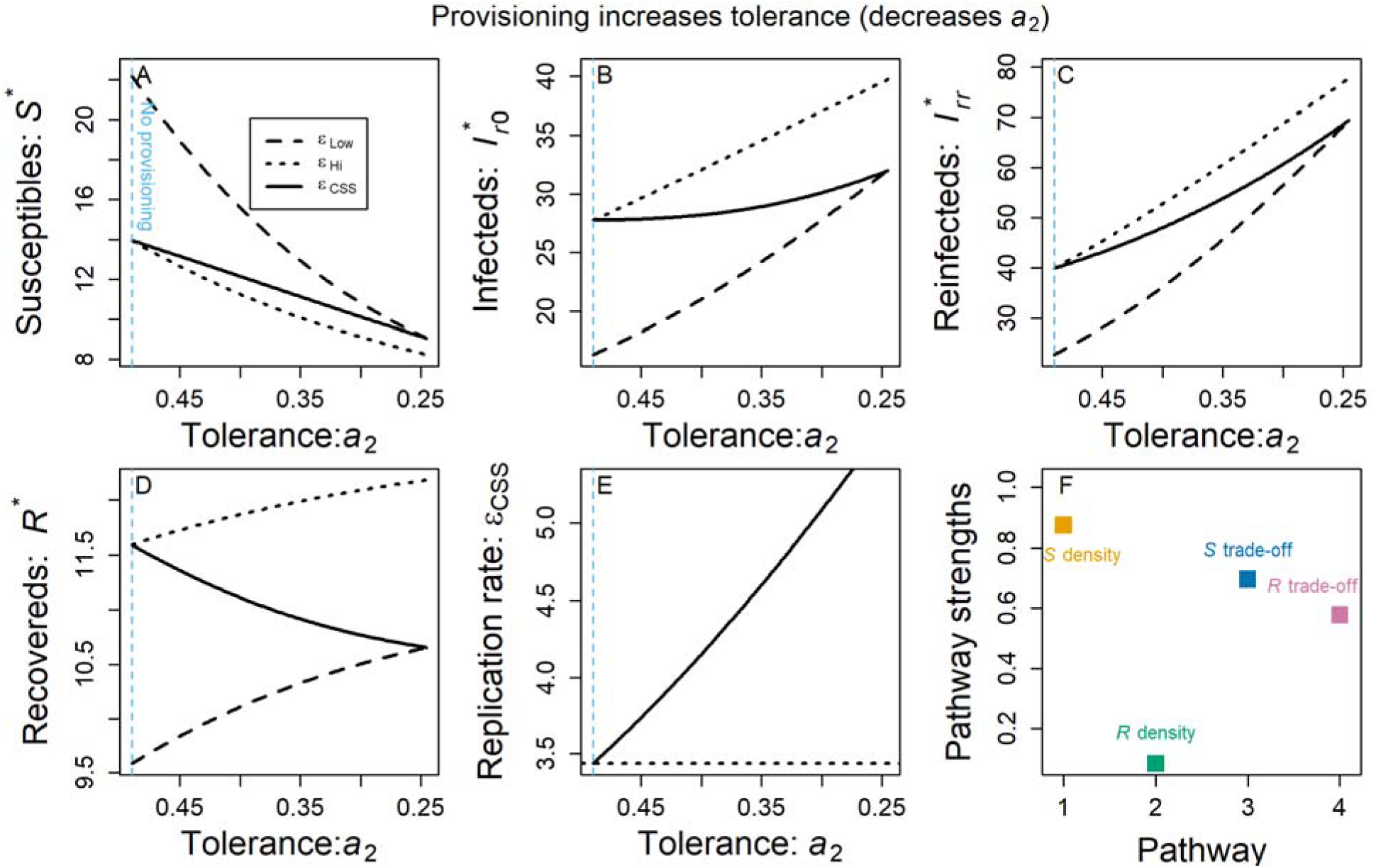
Eco-evolutionary effects of provisioning that strengthens tolerance. We show the effect of stronger tolerance (through lower *a*_2_) and fixed, low replication rate (dotted black), fixed high replication rate (dashed black), and evolved replication rate (solid black) on (A) susceptible host density, (B) infected host density, (C) reinfected host density, (D) and recovered host density. (E) Stronger tolerance strongly selects for higher pathogen replication rate. (F) Positive pathways contribute to selection for higher replication rate while negative pathways contribute to selection for lower replication rate. The sum of pathway strengths gives the net change in *ε*_CSS_ seen across the provisioning range in Fig. A10E. Stronger tolerance drives higher replication rate mainly through positive *S* density, *S* trade-off, and *R* trade-off pathways. The light blue, vertical dashed line shows the parameter value corresponding to default parameter values (i.e., “no provisioning”).

